# Acquisition of hybrid E/M phenotype associated with increased migration, drug resistance and stemness is mediated by reduced miR-18a levels in ER-negative breast cancer

**DOI:** 10.1101/2022.09.05.505398

**Authors:** Madhumathy G Nair, D Apoorva, M Chandrakala, VP Snijesh, CE Anupama, Savitha Rajarajan, Sarthak Sahoo, Gayathri Mohan, Vishnu Sunil Jayakumar, Rakesh S Ramesh, BS Srinath, Mohit Kumar Jolly, Tessy Thomas Maliekal, Jyothi S Prabhu

**Author notes:** Corresponding author- Email id. Equal Contribution (D Apoorva, M Chandrakala). **Ethics approvals and consent to participate:** All procedures performed in the studies involving human participants were in accordance with the ethical standards of both St. John’s Medical College and Hospital (No. 62/2008) and Rangadore Memorial Hospital (RMHEC/02/2010) and with the 1964 Helsinki declaration and its later amendments or comparable ethical standards. A written informed consent for utilization of clinical data was obtained from all enrolled patients. In-vitro experiments were performed at SJRI in accordance with Institutional Biosafety Committee regulations (IBSC/SJRI/01-04/1905/ 2022). Xenograft experiments were performed at the Animal Research Facility, Rajiv Gandhi Centre for Biotechnology (RGCB), Thiruvananthapuram in accordance with ethical standards (AEC No: IAEC/876/TM/2022). **Authors’ contributions:** MGN was involved in design of the work, the acquisition, analysis, and interpretation of data, and has drafted the work; AD and CM have contributed equally towards acquisition, analysis and drafting; SVP and SS have contributed in analysis; ACE and SR has contributed in data acquisition; GM and VSJ has contributed towards acquisition of data (xenograft experiments). RSR and SBS provided clinical samples for the study; TTM, MKJ, and JSP were involved in interpretation of data and drafting of work. All authors have read and approved the final manuscript. All authors give consent for publication.

## Abstract

The complexity of the ER-negative subtype of breast cancer arises due to the heterogeneous nature of the disease rendering them more aggressive and this poses a challenge to effective treatment and eventually the prognosis of the patients. We have explored the miRNA regulation of altered molecular signatures and the effect on tumour progression in ER-negative breast cancer. Using breast tumour specimens, gene expression data from public datasets and in-vitro and in-vivo model systems we have shown that low-levels of miR-18a in ER-negative tumours drives enrichment of hybrid Epithelial/Mesenchymal (E/M) cells with luminal attributes. On inhibition of miR-18a in ER-negative breast cancer cell lines, the cells showed traits of increased migration, stemness and drug-resistance. miR-18a/low tumours were also associated with increased expression of genes associated with EMT, stemness, drug resistance and immune-suppression. Further analysis of the miR-18a targets pointed out at a possible HIF-1α mediated signalling in these tumours. HIF-1α inhibition reduced the enrichment of the hybrid E/M cells and decreased the migratory ability of miR-18a/low cells. Our study reports for the first time a dual role of miR-18a in breast cancer that is subtype specific based on hormone receptor expression and a novel association of low miR-18a levels and enrichment of hybrid E/M cells. The results highlight the possibility of stratifying the ER-negative disease into clinically relevant groups by analysing epigenetic signatures.

## Introduction

Estrogen Receptor (ER) is an important regulator of mammary growth and development. Loss of ER functioning attributed to multiple mechanisms like epigenetic regulation and less dependency on estrogen-based signalling has been linked to endocrine resistance and poor prognosis in ER-positive breast cancer. ER-negative subtype is featured by the absence of ER expression and is associated with a poor prognosis, aggressive disease, and early relapse in comparison to ER-positive breast cancer. Chemotherapy is the main modality of treatment for the ER-negative subtype of breast cancer, however, despite an initial good response there is a high risk of recurrence and distant metastasis in this subtype. The ER-negative subtype is also heterogeneous at the pathological, clinical and at the molecular level (1, 2). The mutational profile of ER-negative breast cancer is vastly distinct from other subtypes. In addition to mutations, epigenetic alterations resulting in deviant gene expression profile are also key contributors to the process of tumour progression in ER-negative breast cancer (3). Deregulated expression of microRNAs (miRNAs) has been attributed to bring about such epigenetic alterations that can affect the process of tumour progression. miRNAs are regulatory RNA molecules that control the process of gene expression, and the aberrant expression of miRNAs are implicated in the process of aggressive disease progression in ER-negative breast cancer. Apart from their role in tumour progression, miRNAs like miR-34 and miR-200 have also been linked to the regulation of plasticity required for cells to transition between the various phases of Epithelial to Mesenchymal Transition (EMT) (4). EMT is a phenomenon characterised by cellular and phenotypic changes that renders plasticity to undergo transformations critical to migration, immune evasion, distant organ seeding and eventually metastasis. This plasticity rendered enables the cells to shuttle between Epithelial, Mesenchymal and the Epithelial-Mesenchymal Hybrid (E/M hybrid) phenotypes and is termed Epithelial-Mesenchymal Plasticity (EMP). The E/M hybrid state consists of cells that are highly tumourigenic and stem cell-like. In addition, they also possess both epithelial and mesenchymal features rendering them abilities for collective cell migration, immune evasion, higher tumour initiating ability which are factors associated with a worse clinical prognosis (5).

miR-18a belongs to the miR-17-92 cluster and has been reported to play a key role in malignant progression of multiple cancer types including lung, gastric, cervical, prostate, breast cancer and osteosarcoma. miR-18a is also reported to have a multifaceted role in tumour progression. It has been reported to promote cancer progression in non-small-cell lung cancer, cervical cancer, and prostate cancer. On the contrary, it has shown to play tumour suppressor roles in pancreatic and colorectal cancer (6, 7). We have previously shown the role of high levels of this miRNA in promoting poor prognosis in ER-positive breast cancer by activating Wnt signalling and bringing about actin remodelling and immune suppression (8, 9). Here, we report for the first time, a dual role of miR-18a in breast cancer that is subtype specific based on hormone receptor expression. We also report a novel association of low miR-18a levels and the enrichment of hybrid E/M cells that leads to phenotypic changes including that of increased migration and stemness in a subgroup of ER-negative tumours.

## Methods

### 1. Cell Lines, culture, and transfection with miR-18a synthetic inhibitors

The cell line MDA-MB-468 was obtained from NCCS (Pune, India, where cell authentication was performed using STR profiling). MDA-MB-468 was cultured in DMEM supplemented with 10% FBS. MDA-MB-231 was obtained from the American Type Culture Collection (ATCC), Manassas, VA. The culture conditions and the phenotypic characterization of the cell line have been reported previously (10). For all experimental assays using cell lines, a passage number below 20 was used and all cell lines were subjected to frequent recharacterization by immunophenotyping and testing of mycoplasma. microOFF™ miRNA inhibitor for miR-18a was purchased from Guangzhou RiboBio Co., Ltd. hsa-miR-18a-5p antagomiR was purchased from Shanghai GenePharma Co., Ltd. The miR-inhibitor/ antagomiR was transfected into cultured MDA-MB-468 and MDA-MB-231 using Lipofectamine RNAiMAX Transfection Reagent according to the manufacturer’s protocol. Briefly, 0.1 × 10^6^ cells were seeded in a 12-well plate in antibiotic free media with 10% FBS. The following day microOFF™ miRNA inhibitor/ Hsa-miR-18a-5p antagomiR (hsa-miR-18a-5p CUAUCUGCACUA GAUGCACCUUA) were mixed with riboFECT™ CP Buffer. A nonspecific microOFF™ inhibitor negative control (cel-miR-239b-5p MIMAT0000295 UUUGUACUACACAAAAGUACUG) or antagomiR negative control was used as the scrambled or negative control. The final concentration of the inhibitor/antagomiR and scrambled was 50-100 nM. To this complex 3 µL Lipofectamine RNAiMAX was added and incubated for 45 min. The transfection complex was then added to the cells along with antibiotic free media with 10% FBS and full distribution over the plate surface was ensured. The cells were incubated for a period of 48-72 h before harvesting. The cells after miR-18a inhibition will be referred to hereafter as MDA-MB-468/miR-18a/inh and MDA-MB-231/miR-18a/inh and the cells transfected with the negative control will be referred to as MDA-MB-468/miR-18a/cont and MDA-MB-231/miR-18a/cont. The transfection efficiency was evaluated by assessing levels of the microRNA targets by blot and q-PCR after 48-72 h. For HIF-1α pathway inhibition, MDA-MB-468/miR-18a/inh cells were treated with a small molecule inhibitor CAY10585, 4 h after transfection. After 72 h, cells were harvested for various assays.

### 2. Protein expression analysis by western blot

The protein expression was assayed and densitometric analysis was performed using quantity one software (Bio-Rad) as reported previously (11). The list of antibodies used is listed in the Supplementary Materials (Supplementary Table S1).

### 3. Immunophenotyping by flow-cytometry

Briefly, cells were trypsinised and post recovery, washed with PBS, fixed in 4% PFA for 10 min followed by permeabilisation in 0.2% Triton X-100 in PBS. Cells were then incubated for 1 h in primary antibodies for CD44 and CD24 at specific dilutions (Supplementary Table S1) and then labelled with specific secondary antibodies. Cells were then re-suspended in 600 μL of PBS and analysed using FACSCalibur cytometer (BD Biosciences). The percentage of CD44^high^ CD24^low^, CD44^low^ CD24^high^, CD44^high^ CD24^high^ expressing cells were analysed. Appropriate secondary antibody controls were included for the analysis. The FL1-H channel was used to detect CD44 and the FL2-H channel was used for detection of CD24.

### 4. Dual immunofluorescence

Cells were grown in poly-L-lysine coated coverslips and transfected as described above. Immunofluorescence was performed by incubating cells in primary antibody anti-E-cadherin and anti-Vimentin overnight at 4°C at specific dilutions (Supplementary Table S1) and then labelled with specific secondary antibodies. The slide was then mounted on gold antifade reagent with DAPI and examined under a fluorescent microscope (Olympus BX51).

### 5. Computational analysis for correlation with E/M hybrid score

The ER-negative tumours of the TCGA-PanCancer Atlas (n=211) and the METABRIC Nature 2012 and Nat Commun 2016 cohorts (n=265) were segregated based on the upper and lower quartiles of miR-18a expression. The TCGA series with n=50 (miR-18a/low) and n=57 (miR-18a/high) tumours and the METABRIC series with n=54 (miR-18a/low) and n=62 (miR-18a/high) tumours were used for further analysis. The clinic-pathological features of the tumours used for the study are enlisted in Supplementary Table S2 and S3. The TCGA data were accessed from the TCGA Research Network: https://www.cancer.gov/tcga (accessed on 15 November 2020), and the METABRIC data were accessed from the European Genome-phenome Archive (12). We used 4 gene signatures to score the individual patient samples to characterize their luminal, basal, epithelial and mesenchymal characteristics. The gene lists for the luminal and basal signature were obtained from a cumulative list of genes (Supplementary Table S4) listed from previously published reports (13-17) and the gene lists for the epithelial and mesenchymal programs were obtained from Tan et al, EMBO Mol. Med. 2014 (18). To calculate the scores we used the ssGSEA algorithm (19) present as a part of gseapy python package.

### 6. Breast cancer tumour specimens used for gene expression analysis

Tumour samples used for molecular analysis were obtained from surgically excised breast tumour specimens from 446 patients enrolled prospectively at two tertiary-care hospitals (St. John’s Medical College and Hospital and Rangadore Memorial Hospital) in Bangalore, from June 2008 to February 2013. Informed consent for use of the material for research was obtained from all patients and the study was approved by the IERB (Institutional Ethics Review Board) at both hospitals (St. John’s Medical College and Hospital (No. 62/2008) and Rangadore Memorial Hospital (RMHEC/02/2010)). Samples were fixed in 10% neutral buffered formalin at room temperature (RT) and stored as formalin-fixed paraffin-embedded (FFPE) blocks. From the set of treatment naive tumour samples (n=275), ER-negative tumour blocks (n=105) and ER-positive tumour blocks (n=170) that met Quality control (QC) criteria for molecular analysis were used for mRNA and miRNA expression analysis. The clinico-pathological features of the ER-negative tumours used for the study are enlisted in Supplementary Table S5.

A set of tissue samples from surgically excised breast tumours with residual disease post neo-adjuvant chemotherapy (NACT) including partial and non-responders (n=54) have also been used. All patients were treated with chemotherapy regimens that included anthracyclines and/or taxanes. Of the 54 residual sections, 43 had adequate tissue for further analysis. Of the 43, 24 qualified for miRNA expression analysis. 34 had sufficient tissue for performing IHC. The clinic-pathological features of the tumours used for the study are enlisted in Supplementary Table S6.

### 7. mRNA and miRNA expression analysis using quantitative PCR

Quantitation of RNA, cDNA synthesis, and q-PCR experiments were performed on tumour specimens and cell line lysates as reported previously (11, 20). The primer sequences for the genes tested are given in Supplementary Table S7. miRNA present in total RNA was extracted and converted to cDNA using Stem-loop primers specific for the chosen miRNA as described previously (8). RU48 was used as endogenous control for normalisation.

### 8. Analysis of mutational spectrum of breast tumours of the METABRIC cohort

The ER-negative tumours of the METABRIC Nature 2012 and Nat Commun 2016 cohorts (n=265) were segregated based on the upper and lower quartiles of miR-18a expression into n=54 (miR-18a/low) and n=62 (miR-18a/high) tumours. Mutational spectrum of cancer driver genes which were collected from IntoGen database was examined (21). The deleterious variants with IMPACT ‘HIGH’ or ‘MODERATE’ were only considered for the analysis. The genes were selected if that gene is mutated at least 3 times across samples. Fisher’s exact test was performed to confirm the significance of the mutation between the miR-18a low and high tumour samples.

### 9. Analysis of Differentially Expressed Genes (DEGs) and pathways in breast tumours of the TCGA and METABRIC series

The ER-negative tumours of the TCGA-PanCancer Atlas (n=211) and the METABRIC Nature 2012 and Nat Commun 2016 cohorts (n=265) were segregated based on the upper and lower quartiles of miR-18a expression as described above. Significant DEGs between miR-18a/high and miR-18a/low groups were filtered based on absolute fold change (FC) ≥ 2 and adjusted p ≤0.05. Gene ontology and pathway analysis of DEGs were performed using the ToppGene suite (22). The deregulated pathways derived from DEGs were visualised using GOplot-r packages (23).

### 10. Correlative analysis of published EMT scores with miR-18a expression in breast tumours of the TCGA and METABRIC series

The TCGA series with n=50 (miR-18a/low) and n=57 (miR-18a/high) tumours and the METABRIC series with n=54 (miR-18a/low) and n=62 (miR-18a/high) tumours were used for this analysis. A pan-cancer EMT Signature derived from patient-tumour data of 11 different cancer types (24) was used for analysing the association with miR-18a. Also, a core-gene list of 130 EMT related genes derived from a meta-analysis of 10 GES datasets was also used for this analysis.

### 11. In-vitro cell migration - wound closure assay

48 h after transfection of cells, the media was replaced with low serum media (0.2% Foetal Bovine Serum) and cells were allowed to rest for 6 h. A wound was generated, and images were captured to mark the initiation time (0 h) and after 48 h. The migratory ability was quantified and normalized by measuring relative gap distance and compared between cells transfected with microOFF™ miRNA inhibitor and negative control.

### 12. Immunohistochemistry to evaluate expression of integrin β3

Primary antibody for integrin β3 was applied for 1 h at room temperature. Sections were further incubated with secondary antibody (DAKO REALTM EnVisionTM) for 20 min as per the kit instructions, followed by development of the colour using DAB (DAKO REALTM EnVisionTM) for 10 min. Appropriate positive and negative controls were run for each batch. Staining patterns of integrin β3 were evaluated by a pathologist (J.S.P). The protein expression analysis was done on post-NACT specimens of patients who had a partial response to chemotherapy where the tumour specimens have more of stromal component and less tumour. Hence immunoreactivity of more than 1% of residual tumour epithelial cells was considered as positive expression for integrin β3.

### 13. Evaluation of drug cytotoxicity using MTT

Cells were seeded (20,000) in 96 well microtiter plates and transfected as described above. After 48 h, medium was removed and replaced with 100 μL of media with paclitaxel at various doses from 10 μM to 200 μM for 48 h.

MDA-MB-231 cells were transfected with ALDH1A1-DsRed2N1 plasmid using Lipofectamine2000 as described previously (25). The stably transfected cells were selected with 100 µg/mL Geneticin (G418) and sorted out in FACS Aria II to enrich CSCs, which were then maintained (20 µg/mL of G418) for experimental purpose. These cells were obtained as a gift from T.T.M. miR-18a was inhibited in these cells and the control (DsRed2N1) cells using microOFF™ inhibitor and inhibitor negative control as described above. After 72 h, medium was removed and replaced with 100 μL of media with paclitaxel at various doses from 10 μM to 200 μM for 48 h. MTT assay was performed as reported previously (11).

### 14. Generation of mammospheres and extreme limiting dilution assay (ELDA) to assess clonogenicity

72 h post transfection, MDA-MB-468/miR-18a/inh and MDA-MB-468/miR-18a/cont were trypsinised and seeded to form spheres using DMEM/F12 media supplemented with 20 ng/mL FGF and EGF along with Insulin-Transferrin supplement in low adherent 12 well plates coated with Poly (2-hydroxyethyl methacrylate). After 5 days, the first-generation spheres were serially propagated and reseeded to form second generation spheres in low adherent 96 well plates by serial dilution. Cells were seeded at a frequency of 1000, 500, 100, 10 and up to 1 cell/well in sextuplicate. After 6 days, the spheres were counted and the sphere forming ability was then calculated using the extreme limiting dilution analysis (ELDA) algorithm as previously described (26).

### 15. Estimate analysis and immune cell identification

The ER-negative tumours of the TCGA-PanCancer Atlas (n=211) and the METABRIC Nature 2012 and Nat Commun 2016 cohorts (n=265) were segregated based on the upper and lower quartiles of miR-18a expression. The TCGA gene expression data was used to infer the stromal and immune scores to predict level of infiltrating stromal and immune cells in the tumours along with the cumulative ESTIMATE score using the ESTIMATE algorithm. The normalized gene expression data with standard annotation files from the TCGA and the METABRIC cohorts were also used for deconvolution of infiltrating immune populations by CIBERSORT algorithm as described previously (9). CIBERSORT was run with the following options: relative and absolute modes together, LM22 signature gene file, 1000 permutations, and quantile normalization disabled. Using the filtered data, the proportions of immune cells in the miR-18a/high and miR-18a/low breast tumours were displayed in the form of a proportion plot. The normalized gene expression data with standard annotation files from the TCGA cohort were also uploaded to the Immune Cell Abundance Identifier (ImmuCellAI), to precisely estimate the infiltration score of 24 immune cell types, including 18 T-cell subsets (27).

### 16. Breast xenograft in-vivo studies

miR-18a was inhibited in MDA-MB-468 using hsa-miR-18a-5p antagomiR and antagomiR negative control as described above. After 72 h, 0.5 × 10^6^ cells from each were harvested and suspended in 50 μL of PBS. Mice were randomly distributed into antagomiR (n=10) and control (n=10) groups. Orthotopic tumours were induced by exposing the fourth (inguinal) mammary fat pad of NSG/ NOD-SCID mice at 6-7 weeks of age and injecting them with cells suspended in 50 μL of Matrigel. On observing palpable tumours, the mice were sacrificed after 21 days, and tumour samples were harvested, and weight measured.

### 17. Histopathological analysis and immunostaining of cryosections

Mouse tumours (n=5 from each group) were formalin-fixed, paraffin embedded and 5 µm sections were cut and Haematoxylin/eosin staining was performed following the standard protocol. Tumours (n=5 from each group) were also fixed in 4% PFA and mounted in OCT compound. 8 μm sections were taken and permeabilized with 0.3% Triton X-100 in PBS and blocked with 3% serum. Primary antibody was added at specific dilutions (Supplementary Table S1) and immunofluorescence performed with anti-E-cadherin and anti-Vimentin antibodies as described above.

### 18. miR-18a target prediction

We analysed the potential targets of miR-18a with 6 different databases and miRNA target prediction tools – miRanda, TargetScan, microT-CDS, PicTar, miRTarBase, miRDB. The miR-18a targets predicted by at least 3 prediction programs were selected for further analysis. The mRNA levels of these targets and their association with miR-18a transcript levels were examined in the miR-18a low and high samples of the TCGA and the METABRIC series.

### 19. Statistical analysis

Descriptive statistics were used for all clinical variables. The difference in gene expression levels was evaluated by the Mann–Whitney U test/Kruskal–Wallis test or the two-tailed Student’s t-test. Correlations were evaluated by Pearsons’s rank test. Kaplan-Meier analysis was used to examine the estimated differences in disease-free survival between the miR-18a/high and miR-18a/low groups. Log-rank test (Mantel-Cox) was used to compare the survival between groups. For in-vitro experimentations, the results are depicted as mean ± standard error of mean calculated from three independent experiments and statistical analysis was performed using Student’s t-test. For all tests, p<0.05 was considered to be statistically significant. All statistical analysis was carried out using the software XLSTAT 2022.2.1.

## Results

### 1. Low levels of miR-18a enriches for the hybrid epithelial/mesenchymal/ lumino-basal phenotype in ER-negative breast cancer

The high levels of miR-18a expression and its effect on prognosis and drug resistance in triple negative breast cancers has been reported previously (28, 29). Evaluation of the levels of miR-18a in 275 breast tumour samples by q-PCR showed that miR-18a was highly expressed (p<0.0001) in the ER-negative tumours (n=105) when compared to ER-positive samples (n=170) (Figure 1A). To further probe the role of miR-18a in ER-negative tumours, miR-18a was inhibited using microOFF™ miRNA inhibitor in breast cancer cell lines. We measured the protein levels of TNFAIP3, an experimentally validated target of miR-18a to assess transfection efficiency. We have previously shown the effective repression of TNFAIP3 protein with miR-18a over-expression (8). In MDA-MB-468/miR-18a/inh cells, we observed a 45% increase in the levels of TNFAIP3 (p=0.0002, Figure 1B). The levels of other targets of miR-18a were assessed for by q-PCR after miR-18a inhibition in both MDA-MB-468 and MDA-MB-231. The levels of miR-18a target genes-*BIRC3, HIF1A, DICER and CDK19* increased in the cell lines after miR-18a inhibition (Supplementary Figure 1). Since miR-18a is involved in epigenetic regulation of the estrogen receptor, we probed for the expression of Keratin 19 that is typically expressed in luminal epithelial cells. On miR-18a inhibition, Keratin 19 levels increased by 50% (p=0.01) in MDA-MB-231 (Figure 1B). This increase in the levels of a luminal cytokeratin in an ER-negative cell line, was intriguing and to examine the possibility of enrichment of luminal-basal hybrid cells, we analysed the ER-negative tumours of the TCGA and the METABRIC cohorts. The miR-18a/low tumours had higher expression of the luminality associated genes like *ESR1, GATA3, FOXA1, XBP1 and TFF1* and a lower expression of the basality associated genes such as *KRT18, KRT17, FOXC1, ANLN, STIL* and *MIA* (p<0.05) (Figure 1C). These tumours were also analysed for the mutational spectrum of the cancer driver genes and this analysis showed a significant mutation load of PIK3CA in mir-18a/low tumours when compared to mir-18a/high tumours (p= 2.15 × 10^−05^ and odds ratio: 8.71 × 10^−02^) (Figure 1D). PIK3CA mutations are most frequently found in ER-positive tumours and has a strong correlation with estrogen receptor signalling. Further, a computational analysis based on gene signatures to score the individual patient samples to characterize their luminal and basal program further supported the hypothesis of enrichment of luminal-basal hybrid cells in miR-18a low tumours. Low levels of miR-18a leads to a less basal and more luminal phenotype in ER-negative tumours in both TCGA and METABRIC tumours (p<0.0001) (Figure 1E). ER-negative tumours of our cohort were also stratified based on miR-18a expression into high (n=24) and low (n=30) groups based on the upper and lower quartiles of miR-18a expression. There was a significantly negative correlation between miR-18a and PGR transcript, an estrogen regulated gene (Pearson’s correlation co-efficient: −0.31, p=0.02) (Figure 1F) in these tumours.

**Figure 1.**
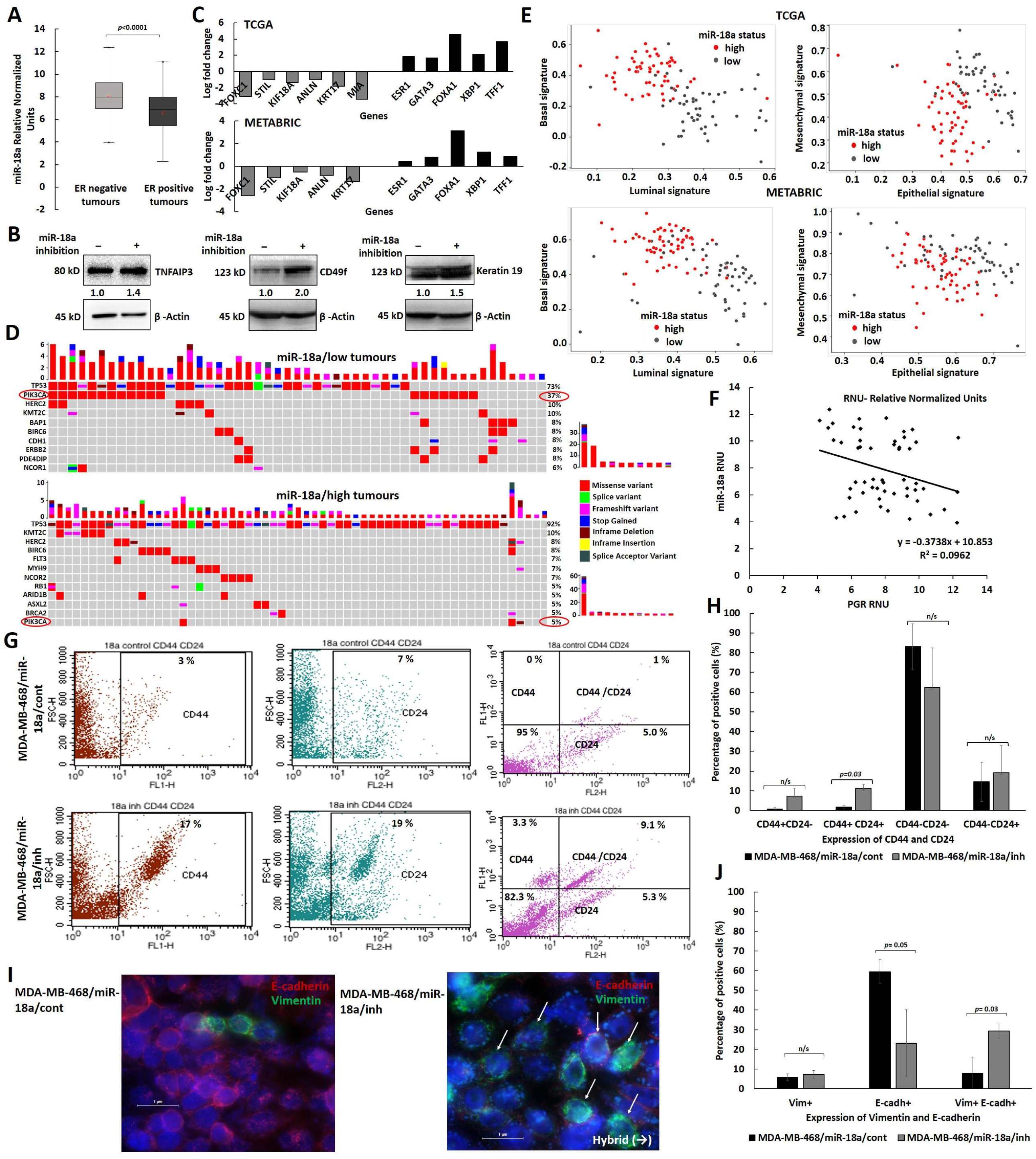
Reduced levels of miR-18a enriches for hybrid EMT/luminal-basal phenotype in ER-negative Breast cancer. (A) Expression levels of miR-18a transcripts in ER-negative (n=105) and ER-positive (n=170) tumours (part of our cohort) as examined by q-PCR. (B) Change in protein expression of TNFAIP3 between MDA-MB-468/miR-18a/cont and MDA-MB-468/miR-18a/inh and Keratin 19, CD49f between MDA-MB-231/miR-18a/cont and MDA-MB-231/miR-18a/inh cells. (C) Expression levels of luminality and basality associated genes in miR-18a/low tumours of the TCGA and METABRIC cohorts when compared to miR-18a/high tumours in the ER-negative subtype (D) Mutational spectrum analysis depicting higher PIK3CA mutation load in miR-18a/low in comparison to miR-18a/high ER-negative tumours of METABRIC cohort. (E) Gene signature based computational analysis characterising luminal, basal, epithelial and mesenchymal scores/signatures in miR-18a/low and miR-18a/high, ER-negative tumours of the TCGA and METABRIC cohorts. (F) Correlation analysis between miR-18a and *PGR* transcript levels in the miR-18a/high (n=24) and miR-18a/low (n=30) ER-negative tumours (part of our cohort) as examined by q-PCR. (G) Representative immunophenotyping images depicting the expression levels of CD44 and CD24 in MDA-MB-468/miR-18a/cont and MDA-MB-468/miR-18a/inh cells as assessed by flow-cytometry. (H) Quantitative assessment of CD44 and CD24 expression in MDA-MB-468/miR-18a/cont and MDA-MB-468/miR-18a/inh as assessed by flow cytometry (cumulative result from 3 independent trials). (I) Representative immunofluorescence images demonstrating the expression of E-cadherin and Vimentin in MDA-MB-468/miR-18a/cont and MDA-MB-468/miR-18a/inh (J) Quantitative assessment of E-cadherin and Vimentin expression in MDA-MB-468/miR-18a/cont and MDA-MB-468/miR-18a/inh.

Cells with hybrid luminal/basal characteristics tend to be enriched for hybrid epithelial/mesenchymal and stemness traits. Hence, we examined the expression of stemness associated protein integrin alpha 6/CD49f in MDA-MB-231/miR-18a/inh cells and the expression doubled on miR-18a inhibition (p=0.0006) (Figure 1B). The cells were also examined for CD44 and CD24 expression as double positivity for CD44 and CD24 is a trait of hybrid E/M cells. The percentage of CD44^+^ CD24^+^ cells significantly increased on miR-18a inhibition by 9% (p=0.03) (Figure 1G & H) in MDA-MB-468. Since another characteristic trait of hybrid epithelial/mesenchymal cells is the dual positivity for Vimentin and E-cadherin, we evaluated the change in expression of these markers in MDA-MB-468/miR-18a/inh cells. There was a significant loss in the expression of E-cadherin (p=0.05), however, the percentage of the dual positive Vimentin^+^ E-cadherin^+^ cells significantly increased after miR-18a inhibition (p=0.03) (Figure 1I & J). This observation was further supported by the computational analysis based on gene signatures on the TCGA and the METABRIC series of tumours. Low miR-18a levels were associated with an increase in both epithelial (p<0.005) and mesenchymal gene signature (p=0.0005) (Figure 1E). These observations indicate that miR-18a low tumours may be enriched for cells with the lumino-basal attributes that overlaps significantly with the hybrid E/M phenotype.

### 2. ER-negative breast cancer with low miR-18a is associated with slower proliferation rates and increased EMT traits

The ER-negative tumour samples of our breast cancer cohort were examined for the association of miR-18a with the Ki67 proliferation index. The tumours were stratified based on miR-18a levels; the tumours with less than the lower quartile expression of miR-18a (miR-18a/low) (n=24) was compared with all the other tumours (miR-18a/high) (n=81). Tumours with Ki-67 index of 14 or more were considered as high proliferative and the tumours with less than 14 were grouped as low proliferative. 67% of the miR-18a/low tumours had a lower Ki67 expression when compared to 24% of the miR-18a/high tumours (p<0.0001) (Figure 2A). The miR-18a levels were further used to correlate with a probability distribution of proliferation score published previously (15) that was derived by fitting a binomial logistic regression model using 2 genes - ANLN and BCL2 as predictor and grade 3 as the determinant. The miR-18a/low tumours were associated with a lower proliferation score (p<0.05) (Figure 2B) when compared to the miR-18a/high tumours. This further suggests that miR-18a/low tumours enriched for hybrid E/M cells may be slow-proliferative and low-cycling than miR-18a/high tumours.

**Figure 2:**
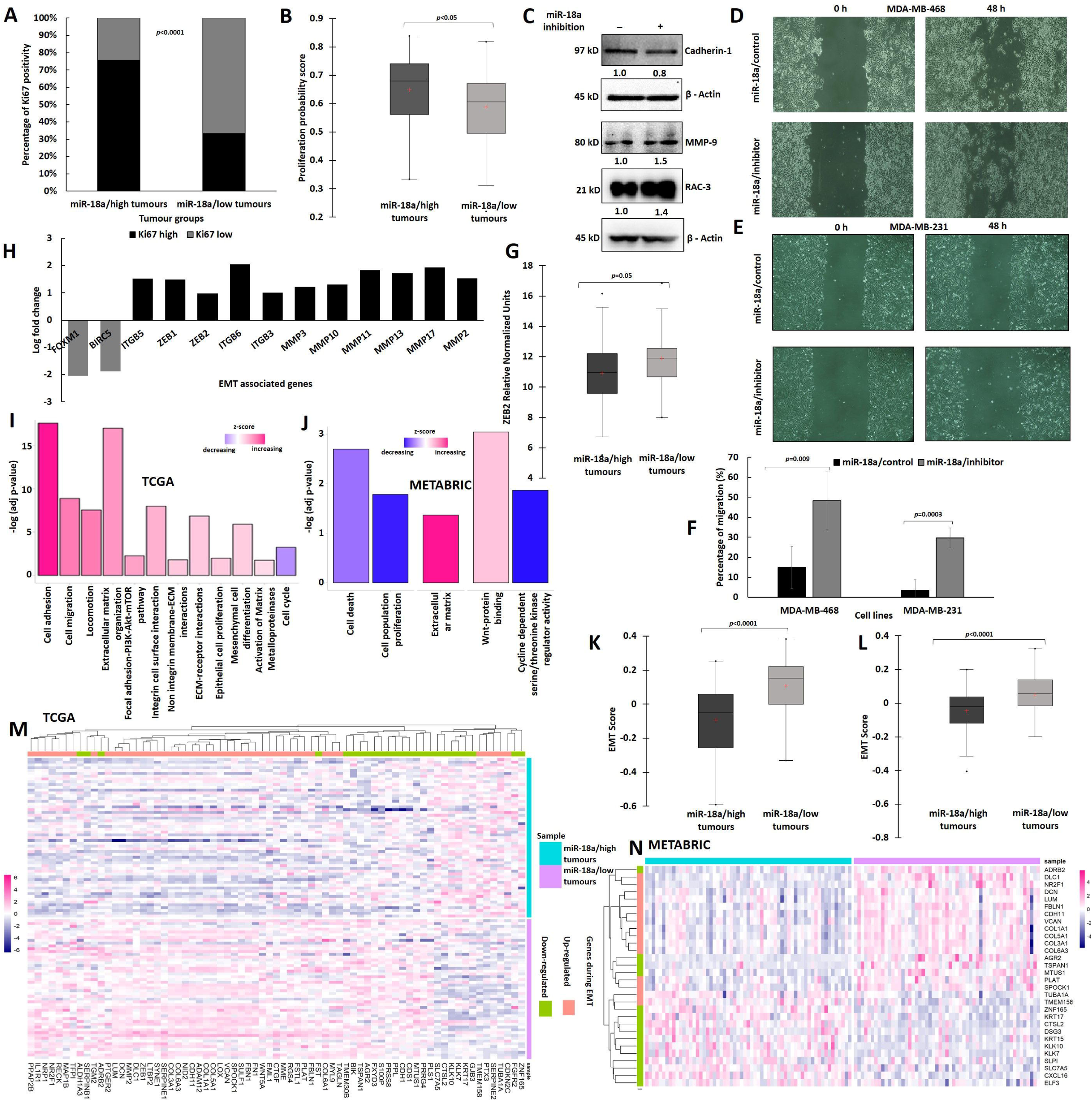
Association of reduced miR-18a levels with proliferation and EMT characteristics in ER-negative breast cancer. (A) Association of Ki67 index and miR-18a levels in miR-18a/high and miR-18a/low tumours & (B) Association of proliferation probability score and miR-18a in miR-18a/high and miR-18a/low tumours of our cohort. (C) Change in protein expression levels of EMT associated proteins in MDA-MB-231/miR-18a/cont and MDA-MB-231/miR-18a/inh cells. (D&E) Migratory ability as assessed by wound healing assay in MDA-MB-468/miR-18a/cont vs. MDA-MB-468/miR-18a/inh and MDA-MB-231/miR-18a/cont vs. MDA-MB-231/miR-18a/inh respectively. (F) Percentage of migration from three independent trials in MDA-MB-468/miR-18a/cont vs. MDA-MB-468/miR-18a/inh and MDA-MB-231/miR-18a/cont vs. MB-231/miR-18a/inh. (G) Association of ZEB2 transcript levels and miR-18a in miR-18a/high and miR-18a/low tumours as assessed by q-PCR (H) DEGs associated with EMT and cell proliferation in miR-18a/low, ER-negative tumours of TCGA (I&J) Functional enrichment of DEGs depicting up-regulated and down-regulated pathways in miR-18a/low, ER-negative tumours of TCGA and METABRIC cohorts respectively. (K) Evaluation of the levels of the EMT score derived from a pan-cancer 77 EMT gene signature in miR-18a/high and miR-18a/low, ER-negative tumours of TCGA and (L) METABRIC cohorts respectively. (M) Heat map depicting expression of genes upregulated and downregulated during the process of EMT (derived from EMT-core list of 130 genes) in miR-18a/low and miR-18a/high groups of ER-negative tumours respectively of TCGA and (N) METABRIC cohorts.

To further probe the role of miR-18a in the process of EMT, the levels of E-cadherin, MMP-9 and RAC3 were probed for in MDA-MB-231/miR-18a/inh cells. There was a significant loss of E-cadherin protein (up to 15%, p=0.03) that is typically lost in epithelial cells to mark the beginning of contact inhibition and increased migration. Rac3 is another critical protein required to regulate adhesiveness and motility in breast cancer. The levels of Rac3 increased by 40% (p=0.0008) and MMP9 levels increased by 55% (p=0.002) (Figure 2C). Further to confirm the increase in migratory ability, a wound healing assay was performed in both MDA-MB-231 and MDA-MB-468 cell lines. After miR-18a inhibition, migratory ability increased by 33% in MDA-MB-468 (p=0.009) and by 26% in MDA-MB-231 (p=0.0003) (Figure 2D, E, F). The observations were further confirmed using the ER-negative breast tumour specimens, (n=105) of our cohort where the association of miR-18a with ZEB2; a master regulator of the EMT process was evaluated. The miR-18a/low tumours segregated based on the lower quartiles of miR-18a expression were found to express high levels of *ZEB2* (p=0.05) (Figure 2G) as assessed by q-PCR. To evaluate these findings in a larger cohort of tumours, the ER-negative tumours of the TCGA and the METABRIC cohort were stratified as miR-18a/low and miR-18a/high as described in the methods. Analysis of the DEGs in TCGA tumours revealed that the miR-18a/low tumours expressed high levels of EMT master regulators-*ZEB1* and *ZEB2* and Matrix metalloproteinases -*MMP2, MMP3, MMP10, MMP11, MMP13* and *MMP17* (p<0.05). The level of *ITGB5*, a glycoprotein involved in facilitating cell migration and angiogenesis was also highly expressed in these tumours (p<0.05) (Figure 2H). METABRIC miR-18a/low tumours displayed elevated levels of *MMP2* (p<0.05) (Supplementary Figure 2). The level of cell proliferation associated genes-*FOXM1* and *BIRC5* genes were expressed less in the miR-18a/low tumours (p<0.05) (Figure 2H). Both FOXM1 and BIRC5 are implicated in driving tumour progression by increasing the cell proliferation rates (30, 31). This result is indeed a reflection of the earlier results observed in our series of tumours where miR-18a/low tumours had a lower Ki67 index. Functional enrichment of differentially expressed genes (DEGs) demonstrated upregulation of the pathways related to cell motility and migration, ECM activation, pathways related to activation of Matrix Metalloproteases, Wnt signalling, and Focal adhesion-PI3K-Akt signalling in TCGA (Figure 2I) and METABRIC series (Figure 2J) of tumours (p<0.05). Moreover, pathways related to cell proliferation and cell cycle were down regulated in these tumours (p<0.05) (Figure 2I & J). Analysis of the tumours using a pan-cancer 77 EMT gene signature derived from 11 cancer subtypes, showed that miR-18a/low tumours of both TCGA and METABRIC series were associated with a higher EMT score (p<0.0001) (Figure 2K, L). Association of miR-18a levels were also examined with a core-gene list of 130 EMT related genes derived from a meta-analysis as described in the methods. In majority of the miR-18a/low tumours, there was a higher expression of the EMT related genes that are upregulated as part of the 130 EMT core-gene list and lower expression of genes that are down-regulated in the EMT core-gene list. These results further confirm our observation that miR-18a/low ER-negative tumours may be slow proliferative but have traits pertaining to increased cell locomotion and migration (Figure 2M, N).

### 3. Low levels of miR-18a are associated with increased chemoresistance and cancer stemness in the ER-negative subtype

Analysis of the genes that are differentially expressed in TCGA miR-18a/low tumours also revealed an increased expression of the genes belonging to the ABC transporter family namely ABCC11, ABCA12, ABCC3, ABCC12, ABCG1, ABCG2 etc (p<0.05) (Figure 3A). ABCC11 and ABCC12 were highly expressed in METABRIC miR-18a/low tumours also (Supplementary Figure 2). Functional enrichment of these DEGs demonstrated upregulation of the pathways related to drug response, drug transporter genes, integrin cell surface interactions, integrin binding and focal-adhesion complex (p<0.05) (Figure 3B). We have shown previously that high integrin β3 levels in triple negative breast cancer contributed to chemoresistance by leading to repression of BAD (11). As integrins are known to trigger downstream pro-survival signalling cascades we looked at the protein expression of integrin β3 by immunohistochemistry in the residual tumours post Neo-adjuvant chemotherapy from partial and non-responders. Integrin β3 protein was expressed by the endothelial cells, stromal immune cells, smooth muscle cells, tumour and peritumoural cells in the residual tumour sections. Figure 3C shows representative IHC images for the expression of integrin β3. The association of miR-18a in integrin β3 negative and positive tumours were probed for in ER-negative residual tumours (n=13). miR-18a/low tumours expressed higher levels of integrin β3 (p=0.1) when compared to miR-18a/high tumours (Figure 3D). To further confirm the findings, drug resistance was measured after inhibition of miR-18a in MDA-MB-468 with paclitaxel at varying doses from 10 µM to 200 µM. The drug sensitivity of MDA-MB-468/miR-18a/inh was not different from MDA-MB-468/miR-18a/cont at various doses from 10 µM to 100 µM (p>0.05). With 200 µM paclitaxel treatment, there was a 12% increase in the cell viability of MDA-MB-468/miR-18a/inh when compared to MDA-MB-468/miR-18a/cont (p=0.005) (Figure 3E).

**Figure 3:**
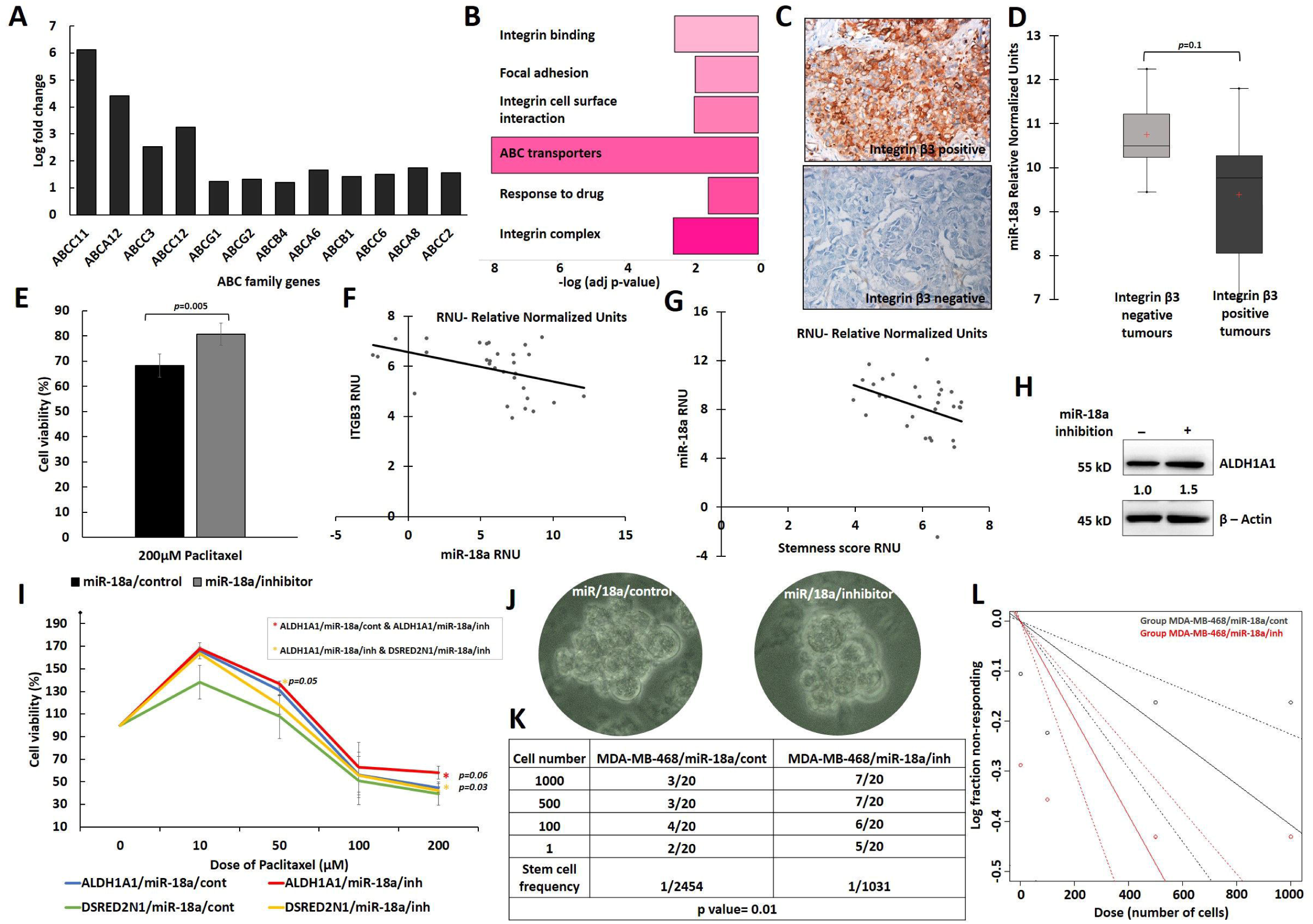
Association of low miR-18a levels with response to drug and stemness acquisition in ER-negative breast cancer. (A) Fold change of DEGs associated with drug transporter gene family (ATP-binding cassette transporter) in miR-18a/low, ER-negative tumours of TCGA cohort. (B) Functional enrichment of DEGs depicting up-regulated pathways in miR-18a/low, ER-negative tumours of TCGA. (C) IHC representative images of Integrin β3 stained sections of residual tumours post NACT (D) Association of Integrin β3 protein and miR-18a in ER-negative residual tumours post NACT (E) Cell viability of MDA-MB-468/miR-18a/cont and MDA-MB-468/miR-18a/inh after 200 µM paclitaxel treatment. (F) Association of *ITGB3* and miR-18a transcript levels; and (G) association of stemness score and miR-18a transcript levels in miR-18a/low treatment naïve primary ER-negative tumours as assessed by q-PCR. (H) Change in expression levels of ALDH1A1 between MDA-MB-468/miR-18a/cont and MDA-MB-468/miR-18a/inh. (I) Percentage of cell viability of ALDH1A1/miR-18a/cont, ALDH1A1/miR-18a/inh, DsRed2N1/miR-18a/cont and DsRed2N1/miR-18a/inh groups in MDA-MB-231 after paclitaxel treatment across doses ranging from 10-200 µM. (J) Representative images of spheres obtained from MDA-MB-468/miR-18a/cont and MDA-MB-468/miR-18a/inh cells (K) Table depicting clonogenicity (stem cell frequency) calculated from extreme limiting dilution assay and (L) graph depicting clonogenicity from extreme limiting dilution assay performed on MDA-MB-468/miR-18a/cont and MDA-MB-468/miR-18a/inh groups across the same initial seeding dose range of 1-1000 cells.

Cells with hybrid luminal/basal-epithelial/mesenchymal features tend to display enhanced cancer stem cell (CSC) properties. Integrin β3/CD61 is also identified as a mammary progenitor marker that identifies the cancer stem cell population enriched for tumourigenic potential (32). The association of miR-18a/low tumours with integrin β3 in post-neoadjuvant residual tumours intrigued us to probe for the same association in primary treatment naïve tumours of our cohort. ER-negative tumours of our cohort were also stratified based on miR-18a lower quartile expression into miR-18a/low (n=30) tumours. Within miR-18a/low tumours, there was a significantly negative correlation between *ITGB3* and miR-18a (Pearson’s correlation co-efficient: −0.42, p=0.02) (Figure 3F). To examine other features of cancer stemness displayed by miR-18a/low tumours, q-PCR assay for cancer stemness associated genes – *SALL4, LGR5, BMPR1B* was performed in these tumours. A stemness score was arrived at by calculating the mean gene score of *SALL4, LGR5, BMPR1B and ITGB3*. Within miR-18a low tumours, there was a negative association between miR-18a and the stemness score (Pearson’s correlation co-efficient: −0.33, p=0.06) (Figure 3G), implying that as miR-18a levels reduced the stemness score was higher in these ER-negative tumours. The levels of ALDH1A1 and BMP4 was also high in the METABRIC miR-18a/low tumours on analysis of the DEGs (p<0.05) (Supplementary Figure 2). BMP4 is implicated in promoting metastasis in breast cancer by enhancing cancer stemness. On miR-18a inhibition in MDA-MB-468, we also observed an increase in the levels of ALDH1A1 by 50% (p=0.04) (Figure 3H). To further examine the effects of miR-18a inhibition on cells enriched for cancer stemness, MDA-MB-231 cells transfected with ALDH1A1-DsRed2N1 plasmid was used. miR-18a was inhibited in MDA-MB-231 cells with ALDH1A1-DsRed2N1 reporter and the control (DsRed2N1) cells using microOFF™ inhibitor and inhibitor negative control as described in the methods. To confirm the stemness of ALDH1A1-DsRed2N1 cells, the level of stemness associate genes LGR-5 and SOX2 were assessed by q-PCR and were found to be higher than control (DsRed2N1) cells (Supplementary Figure 3). Post transfection, the cells were subjected to drug sensitivity assays with paclitaxel with varying doses from 10 µM to 200 µM. ALDH1A1/miR-18a/inh cells were more resistant (by 14%) than ALDH1A1/miR-18a/cont only at 200 µM dose of paclitaxel (p=0.06). However, ALDH1A1/miR-18a/inh cells were significantly more chemo resistant than DSRED2N1/miR-18a/inh at both 50 µM (by 18%; p=0.05) and 200 µM (by 16%; p=0.03) doses of paclitaxel (Figure 3I). To further confirm the role of low miR-18a in rendering stemness attribute to breast cancer cells, mammosphere forming ability was evaluated in MDA-MB-468/miR-18a/inh cells. MDA-MB-468-miR-18a-cont formed lesser number of distinct spheres than MDA-MB-468-miR-18a-inh (Figure 3J). The spheres were serially propagated, and extreme limiting dilution assay was performed to assess clonogenicity of the spheres formed. The clonogenicity (1/stem cell frequency) was 1/1031 in MDA-MB-468-miR-18a-inh cells and almost two times lower in MDA-MB-468-miR-18a-cont (1/2454) cells (p=0.01) (Figure 3K & L). These results clearly ascertain the effect of diminished miR-18a levels in enhancing cancer stemness attributes and drug resistance in breast cancer cells.

### 4. Decreased levels of miR-18a in ER-negative breast cancer correlates with increased stromal-immune infiltration and immunosuppression

A bioinformatic approach was followed to estimate the proportion of immune infiltrate in miR-18a/low tumours of the TCGA and the METABRIC cohort as described in the methods. The ESTIMATE (Estimation of Stromal and Immune cells in Malignant Tumour tissues using Expression data) is a tool that uses gene expression data for predicting tumour purity, and the presence of infiltrating stromal/immune cells in tumour tissues. ESTIMATE score generates 3 different scores. The stromal score depicts the stromal presence in these tumours and the immune score captures the immune infiltration in the tumours. These two scores form the basis of the ESTIMATE score that is an inference of the tumour purity. The miR-18a/low tumours were associated with a higher stromal (p<0.0001) (Figure 4A) and immune score (p=0.008) (Figure 4B). These tumours also had a higher ESTIMATE score when compared to miR-18a/high tumours (p<0.0001) (Figure 4C).

**Figure 4:**
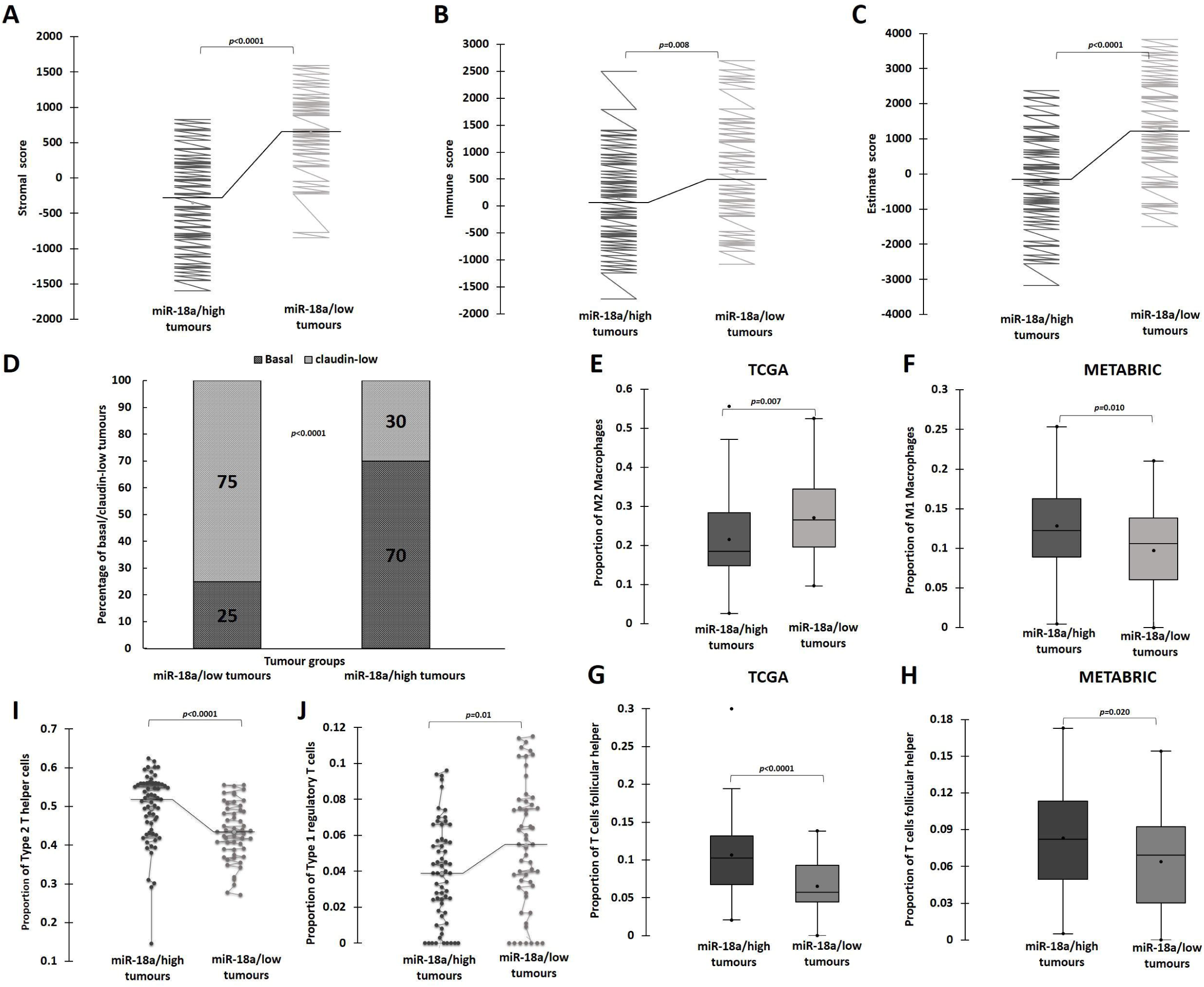
Correlation of low miR-18a levels with immune modulation in ER-negative breast cancer. (A B & C) ESTIMATE generated Stromal score, Immune score and Estimate score for miR-18a/high and miR-18a/low tumours of TCGA cohort. (D) Graph representing proportion of Claudin-low and Basal tumour subtypes in miR-18a/low and miR-18a/high, ER-negative tumours of METABRIC series. (E&F) CIBERSORT analysis depicting the proportions of M2 and M1 macrophages in miR-18a/high and miR-18a/low tumours (G&H) CIBERSORT analysis depicting the proportions of T follicular helper cells in miR-18a/high and miR-18a/low, ER-negative tumours of TCGA and METABRIC cohorts respectively. (I&J) ImmuneCellAI analysis depicting the proportions of Th2 (Type 2 helper cells) and Tr1 (Type 1 regulatory T cells) in miR-18a/high and miR-18a/low, ER-negative tumours of TCGA series.

Most of the traits we have identified as associated with miR-18a/low tumours overlap with that of the claudin-low tumours. Claudin low subtype represent ER-negative tumours that express EMT genes, and stem-like gene expression patterns. They are also known to have marked stromal and immune infiltration. They are associated with a lower proliferation rate and a lower Ki-67 index when compared to non-claudin low tumours (33). Tumours from the METABRIC series were used to examine the proportion of claudin low and basal tumours in the miR-18a high and low tumours. When only basal and claudin low tumours are grouped together, 21/28 of miR-18a/low tumours represent the claudin-low subtype when compared to only 18/60 of miR-18a/high tumours (p<0.0001) (Figure 4D). Further, to examine if the immune-stromal infiltration in miR-18a/low tumours were suggestive of an immunosuppressive microenvironment, immune cell identification was done using CIBERSORT analysis, a method that describes the cell composition of complex tissue from their gene expression profiles in tumours. The analysis in both the cohorts revealed that ER-negative tumours with low miR-18a correlated with increased proportions of M2 macrophages (TCGA-p=0.007) and decreased proportion of M1 macrophages (METABRIC-p=0.01) (Figure 4E & F). miR-18a/low tumours also had a significantly lower presence of T-follicular helper cells (TCGA-p<0.0001, METABRIC-p=0.02), which are specialised T cells that play a crucial role in protective immunity by helping B cells (34) (Figure 4G & H). Further evaluation of the immune composition was done using ImmuCellAI, a gene expression-based method for estimating the abundance of multiple types of T-cell subsets, in the TCGA cohort. miR-18a/low tumours had a lower proportion of Th2 (Type 2 helper cells) (p<0.0001) and a higher proportion of Tr1 (Type 1 regulatory T cells) (p=0.01) (Figure 4I & J). Th2 cells participate in building anti-tumour immunity and aids in tumour clearance and Tr1 mediate immune suppression and establish peripheral tolerance. Thus, the results are suggestive of an enhanced stromal-immune infiltrate in miR-18a/low tumours that may be pro-tumour and immune-suppressive.

### 5. miR-18a inhibition leads to increased tumour size and enriches for hybrid E/M cells in-vivo. HIF-1α inhibition leads to a reversal of hybrid E/M phenotype in miR-18a inhibited cells

To further decipher the clinical relevance and the prognostic implication of miR-18a low levels in ER-negative breast cancer, a Kaplan Meier survival analysis was performed on the METABRIC series of samples. Tumours were stratified based on median expression levels of miR-18a in both ER-negative (median-8.2) and ER-positive tumour samples (median-7.3). The analysis was then performed between ER-positive/miR-18a/low (n=370), ER-positive/miR-18a/high (n=377), ER-negative/miR-18a/low (n=105) and ER-negative/miR-18a/high (n=106) tumour samples (Figure 5A). There was no significant difference in disease-free survival between the ER-negative/miR-18a/low tumours and the ER-negative/miR-18a/high tumours (p=0.9). However, the disease-free survival was significantly different between the ER-positive/miR-18a/low tumours vs ER-negative/miR-18a/low tumours (p=0.001) and the ER-positive/miR-18a/high tumours vs ER-negative/miR-18a/low tumours (p=0.001). The mean survival time for ER-negative/miR-18a/low tumours was 114 months when compared to 141 months in ER-positive/miR-18a/low tumours. A target scan analysis was performed to decipher the gene targets of miR-18a that may be differentially expressed and driving the effects brought about by the low miR-18a levels in the miR-18a/low tumours of the ER-negative breast subtype. 15 targets identified by at least 3 different target mining softwares were shortlisted (Table 1). The expression levels of these targets were analysed in the tumours of the TCGA by stratifying both ER-positive and ER-negative tumours into 2 groups each based on upper and lower quartiles of miR-18a expression. Of all the targets, *PDE4D* and more significantly, *HIF1A* showed high expression in ER-negative miR-18a/low tumours (Figure 5B). *HIF1A* emerged as the target predicted by all target prediction tools and miR-18a dependent HIF-1α and hypoxic regulation has already been reported in basal breast cancer previously (28). Hence, the levels of HIF-1α protein were probed for in MDA-MB-468/miR-18a/inh cells and the expression doubled post miR-18a inhibition (p=0.05) (Figure 5C). This increase in the HIF-1α levels prompted us to examine the presence of an activated hypoxic gene expression if any. Genes involved in hypoxia were retrieved using literature mining (35-37). The collected genes were mapped to TCGA dataset. Further, the genes which were showing a significant difference (p<0.05) between miR-18a/high and low tumours were filtered. The heatmap representing the pattern of expression of filtered hypoxia genes shows that the hypoxia related genes were upregulated in the miR-18a/low tumours (Supplementary figure 4).

**Table 1.** List of miR-18a targets: Target scan analysis was performed to decipher the gene targets of miR-18a. 15 targets identified by at least 3 different target mining softwares and their known function are enlisted below.

| Sl No. | Gene | Function (Source - GeneCards, NCBI Gene) |
| --- | --- | --- |
| 1 | *HIF1A* | Mediates hypoxia-induced expression of mRNA-encoding genes, regulates the expression of non-coding RNAs, which are critical regulators of migration, invasion, and metastasis. |
| 2 | *BBX* | Transcription factor necessary for cell cycle progression from G1 to S phase |
| 3 | *CDK19* | Mediator kinases, transcriptional co-regulators |
| 4 | *DICER1* | Responsible for cleaving double stranded RNAs into small interfering RNAs and microRNAs |
| 5 | *NEDD9* | Positive regulator of epithelial-mesenchymal transition and promotes invasion |
| 6 | *EPB41L1* | Role in cell adhesion and migration, malignant progression |
| 7 | *ESR1* | Regulates the transcription of estrogen-inducible genes that play a role in growth, metabolism, sexual development, gestation |
| 8 | *GLRB* | Down-regulation of neuronal excitability, generation of inhibitory postsynaptic currents |
| 9 | *INADL* | Mediate protein-protein interactions, regulate the formation and stabilization of tight junctions |
| 10 | *MAP3K1* | Serine/threonine kinase in multiple cell signalling cascades |
| 11 | *PDE4D* | Major regulators of cAMP-hydrolyzing activity |
| 12 | *PHC3* | Transcriptional repression, chromatin remodelling and modification of histones |
| 13 | *RORA* | Interacts with NM23-2, a nucleoside diphosphate kinase involved in organogenesis and differentiation, as well as with NM23-1, the product of a tumour metastasis suppressor candidate gene |
| 14 | *SH3BP4* | Involved in cargo-specific control of clathrin-mediated endocytosis, specifically controlling the internalization of a specific protein receptor. |
| 15 | *ZNF367* | Transcriptionally activates KIF15 and regulates cell cycle |

**Figure 5:**
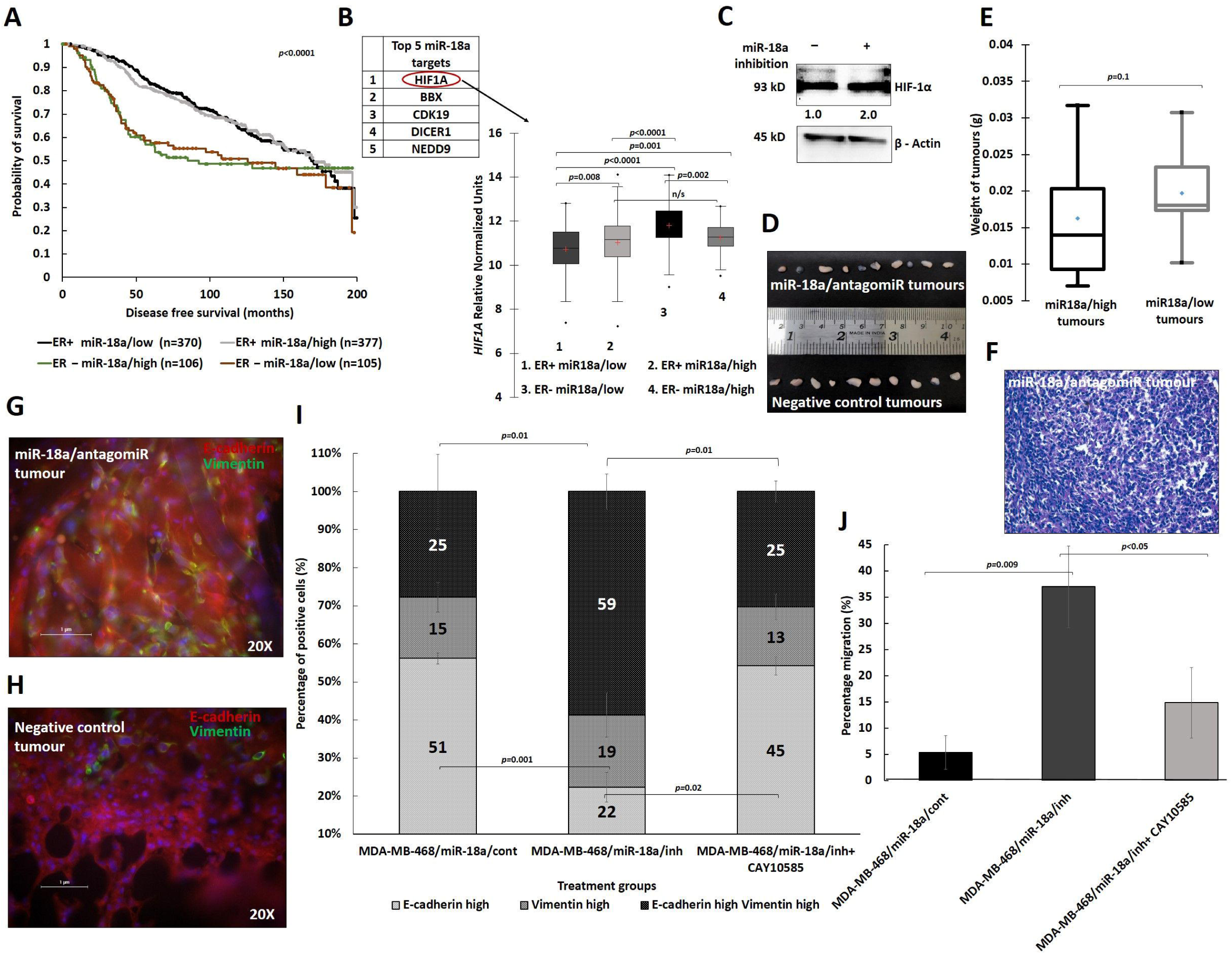
Effect of miR-18a inhibition on tumours in-vivo and association with *HIF1A* expression. (A) Kaplan Meier analysis depicting disease free survival in ER-positive/miR-18a/low, ER-positive/miR-18a/high, ER-negative/miR-18a/low and ER-negative/miR-18a/high tumours of the METABRIC cohort. B) Expression of *HIF1A* in ER-positive/miR-18a/low, ER-positive/miR-18a/high, ER-negative/miR-18a/low and ER-negative/miR-18a/high tumours of the TCGA cohort. (C) Increased expression levels of HIF-1α in MDA-MB-468/miR-18a/inh in comparison to MDA-MB-468/miR-18a/cont. (D) Images of tumours harvested from mice injected with miR-18a/antagomiR cells and antagomiR negative control cells post 21 days of injection (E) Box plot depicting the average weight of the tumours harvested from miR-18a/antagomiR and the antagomiR negative control mice. (F) Representative H&E staining image of tumour from miR-18a/antagomiR mice. (G) Immunofluorescence to demonstrate the dual positivity of E-cadherin and Vimentin in tumour sections from miR-18a/antagomiR tumour and (H) absence of dual positivity of E-cadherin and Vimentin in antagomiR negative control tumour. (I) Quantitative analysis of immunofluorescence for expression of Vimentin and E-cadherin in MDA-MB-468/miR-18a/cont, MDA-MB-468/miR-18a/inh, and MDA-MB-468/miR-18a/inh + CAY10585 (J) Percentage of migration as measured by wound healing assay in MDA-MB-468/miR-18a/cont, MDA-MB-468/miR-18a/inh, and MDA-MB-468/miR-18a/inh + CAY10585.

To further evaluate the in-vivo tumourigenic potential of miR-18a inhibited cells, mouse xenograft experiments were performed. miR-18a was inhibited using miR-18a-5p antagomiR and antagomiR negative control in MDA-MB-231 that were injected to induce orthotopic tumours in NSG/ NOD-SCID mice. The tumours formed with miR-18a/antagomiR cells were larger in volume by 21% when compared to tumours formed from antagomiR negative control cells (p=0.1) (Figure 5D,E). Figure 5F depicts a representative H&E image of a miR-18a/antagomiR tumour. Immunofluorescence staining was performed on the tumour sections from the miR-18a/antagomiR tumours and the antagomiR negative control tumours for the markers Vimentin and E-cadherin. miR-18a/antagomiR tumours showed a higher proportion of cells co-expressing Vimentin and E-cadherin (Figure 5G) when compared to negative control tumours (Figure 5H). The results further support the hypothesis that low levels of miR-18a drives tumour progression by enriching for hybrid E/M cells in ER-negative breast cancer.

Further to confirm the role of HIF-1α in miR-18a inhibited cells, HIF-1α pathway was blocked using CAY10585, a HIF-1α inhibitor that suppresses transcription of HIF-1α target genes. We observed that the percentage of hybrid E/M cells (Vimentin^+^ E-cadherin^+^) reduced by 30% (p=0.01) accompanied by an increase in the proportion of cells that were Vimentin^-^ E-cadherin^+^ (p=0.02) (Figure 5I). Moreover, the migratory ability of the MDA-MB-468/miR-18a/inh cells were found to decrease by 22% with HIF-1α pathway inhibition (p<0.05) (Figure 5J). The results show the existence of a possible HIF-1α dependent pathway regulated by low-levels of miR-18a in the ER-negative subtype driving the hybrid E/M phenotype.

## Discussion

The complexity of the ER-negative subtype of breast cancer arises due to the heterogeneous nature of the disease and this poses a challenge to effective treatment and eventually the prognosis of the patients. The ER-negative subtype is usually more aggressive and has a worse prognosis than the ER-positive subtype. Better understanding of the disease exists with advancement in genomics, however the ER-negative subtype has very few targeted and tailored therapy options. The molecular subtyping studies especially the PAM50 classification has unravelled the existence of a basal-like and Her2-enriched subclasses within the ER-negative subtype. Genome sequencing and mutational profiling done by the TCGA network and multiple other groups have led to identification of mutational signatures that not only affect genomic signatures but also bring about epigenetic changes (1-3). This may be causal for tumour heterogeneity leading to differential activation of signalling pathways that eventually leads to divergent molecular signatures. This contributes towards the aggressive biology and differential response to treatment and further the prognosis. However, there still exists dearth of knowledge on the role played by such epigenetic changes especially miRNA regulation of tumour progression. The miRNA mediated regulation of transcriptomics becomes even more intriguing owing to the multifaceted role of these regulatory molecules that are now being exhumed. More reports emerge with evidence of a single miRNA playing tumour promoting roles and tumour suppressive roles among cancer subtypes. We have previously reported the tumour-promoting role of high levels of miR-18a that leads to Wnt pathway activation thus promoting metastasis and poor prognosis in ER-positive breast cancer. We have also identified that miR-18a driven Wnt pathway activation may be the basis for the ‘immune cold’ phenotype that is displayed by ER-positive tumours (8, 9). In this manuscript we present evidence for an atypical biology that may be driven by low levels of miR-18a in the absence of expression of the hormone receptors-ER and PR.

The ER-negative tumours of our cohort expressed higher levels of miR-18a when compared to ER-positive tumours. Nevertheless, the tumours that expressed lower levels of this miRNA within the ER-negative subtype were found to retain a different biology. These tumours were found to have both epithelial and mesenchymal traits thus exhibiting traits of hybrid E/ M tumours. In addition, they were found to have increased luminal traits also making them luminal/basal hybrid tumours. In-vitro and computational analysis further confirmed these findings. The miR-18a low tumours are associated with a lower Ki-67 index, lower proliferation rates and features of increased migration and EMT. ALDH1A1 levels increased post inhibition of miR-18a and this increased the drug-resistance. Inhibition of miR-18a also increased clonogenicity and mammosphere forming ability. In-silico analysis also showed a correlation of low miR-18a levels to immunosuppression in ER-negative tumours. It was also intriguing that a large proportion of miR-18a/low tumours overlapped with the claudin-low tumours. These findings are clinically relevant as the claudin-low is a much uncharacterised subtype of breast cancer. We did not observe any difference between the prognosis of miR-18a/low and high tumours of ER-negative subtype. This may be attributed towards the heterogeneity in gene expression patterns and mutational profiles generally noted in ER-negative tumours. However, in in-vivo models, miR-18a inhibited cells formed larger tumours and they expressed more hybrid E/M cells. Regulation of a miR-18a hypoxic gene signature by activation of HIF-1α in basal breast cancer has been reported earlier (28). We also observed that the miR-18a/low tumours expressed high levels of *HIF1A*. HIF-1α inhibition using a small molecule in miR-18a inhibited cells brought down the proportion of hybrid E/M cells and the migratory ability. The results mirror the observations reported previously where it was noted that the maintenance of highly tumourigenic E/M hybrid state was brought about by activation of EMT-inducing transcription factors and canonical Wnt signalling in basal breast cancer cells. The role of hypoxia driven signalling through activation of P4HA2 in maintaining the partial or hybrid E/M phenotype in breast cancer was also recently examined (38, 39).

miR-18a regulates ER signalling by binding to the 3’UTR and regulating it’s expression and this may be one of the ways by which epigenetic silencing of ER is mediated during evolution of ER-negative tumours. Higher levels of miR-18a in ER-negative tumours may be an implication of such regulation. However, lower levels of miR-18a in these tumours may be activating pathways required for the enrichment of hybrid E/M cells that leads to multitude of phenotype changes. Recent studies render functionalities of gene expression tuning and expression buffering to miRNAs. Expression buffering is a process by which weakening in the variance of the expression level of the target genes is mediated by miRNAs (40). The biology seen in miR-18a/low tumours may be attributed to such a buffering function where low levels fail to buffer the mean levels of miR-18a target genes such as *HIF1A* and thus lead to alternate phenotypic changes. The blocking of HIF-1α and reversal of hybrid E/M is a confirmation of such altered buffering.

Limitations of this study include a smaller sample size of miR-18a low ER-negative tumours used for the analysis. However, the fact that we were able to verify the observations in two larger datasets increases the credibility of the observations made in clinical specimens. In-vitro results further validate and strengthen our results and observations. The immune suppression observed was identified using only an in-silico approach. Further in-vitro validations need to be performed to confirm the immunosuppressive effects of miR-18a in ER-negative tumours. Since the discovery of miRNAs in the field of cancer, there also has been increase in the focus on the therapeutic implications of these small molecules. The pleiotropic and multifaceted nature of these small regulatory molecules make them attractive drug targets and amenable to tweaking, especially for ER-negative subtype that is vastly heterogeneous. Stratifying these tumours based on epigenetic phenotypic alterations can become a promising strategy for personalised medicine.

## Acknowledgement

We thank the Department of Health Research, Ministry of Health & Family Welfare, and ICMR India, for the Young Scientist fellowship to M.G.N. We are grateful to Nadathur Estates and the Bagaria Education Trust for their support of all the breast cancer research activities at SJRI since 2008. Acknowledgement to pathologist Dr. Sharada Patil for help with histological analysis, Ms. Mahalaxmi S for technical help, Mr. Santhosh HS for research assistance, Ms. Amrita Mohan for help with xenograft experiments at R.G.C.B. Gratitude to Dr. Harikrishna Nakshatri, Indiana University School of Medicine for his inputs. We also thank Dr. Sridhar TS for critical review of the manuscript.

---

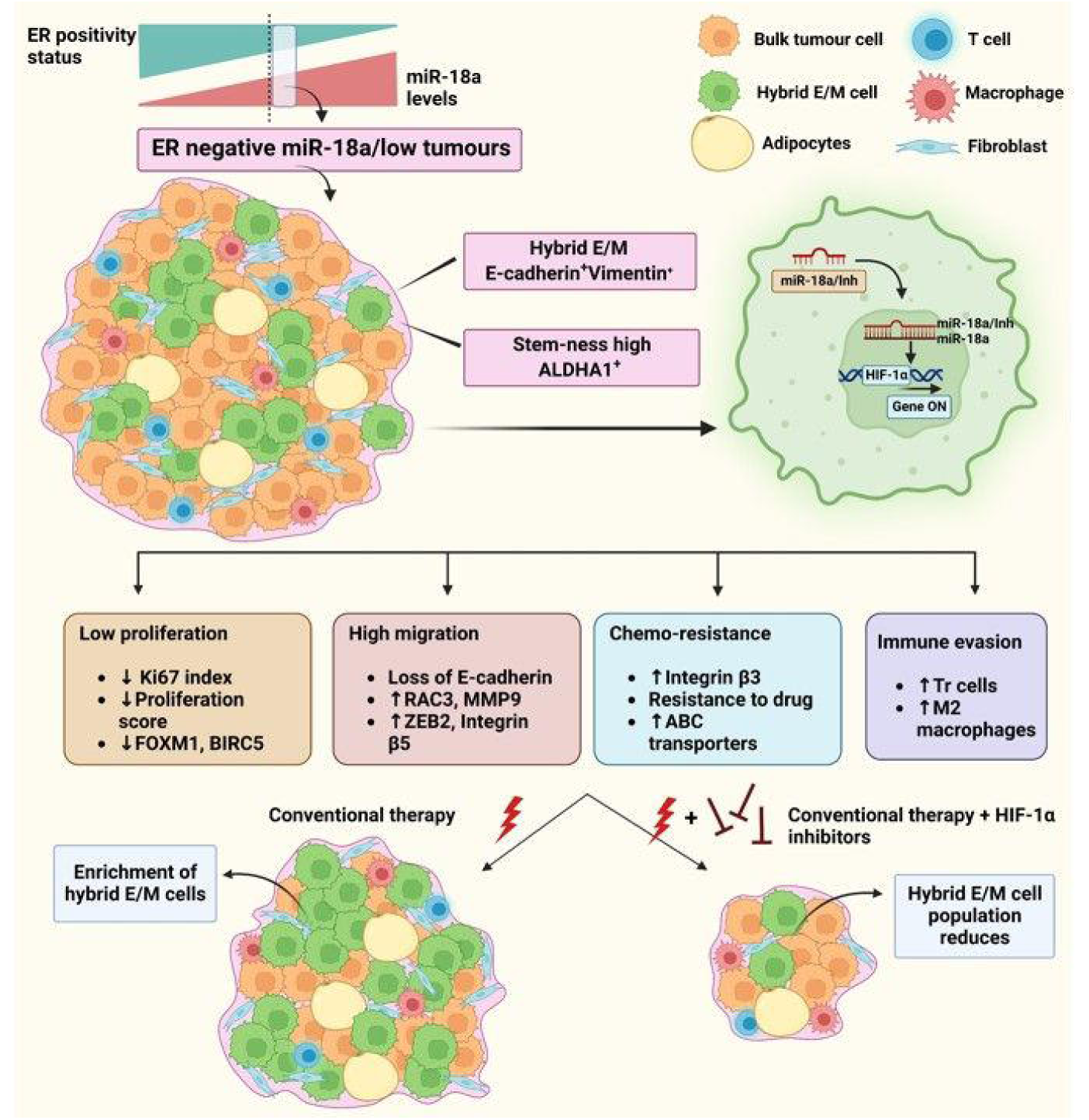
miR-18a propelled hybrid E/M phenotype in ER-negative tumors-. miR-18a is highly expressed by the ER-negative tumors in comparison TO the ER-positive tumors. miR-18a represses ER expression by binding to the 3’UTR which leads to epigenetic silencing of ER leading to evolution of ER-negative tumors. Lower levels of miR-18a in a fraction of these ER-negative tumors may be activating pathways required for the enrichment of hybrid E/M cells. This may be accredited to a faulty buffering function where low levels fail to buffer the mean levels of miR-18a target genes such as *HIFlA* thus leading to an alternate phenotype. This altered phenotype encompasses slow cell cycling, decreased proliferation, increased migration, drug resistance and immune evasion. Conventional therapy regimens may target the bulk of the cells but may lead to enrichment of hybrid E/M cells that are drug resistant. These residual hybrid E/M cells owing to the increased migratory and immune evasive abilities may lead to distant metastasis and recurrence. A combinatorial treatment strategy with HIF-lα inhibitors may effectively eliminate these cells with hybrid E/M attributes.

