## Supplementary data for "Acquisition of hybrid E/M phenotype associated with increased migration, drug resistance and stemness is mediated by reduced miR-18a levels in ER-negative breast cancer"

Supplementary Figure 1: Expression level of *BIRC3*, *HIF1A*, *DICER* and *CDK19* after miR-18a inhibition as measured by q-PCR

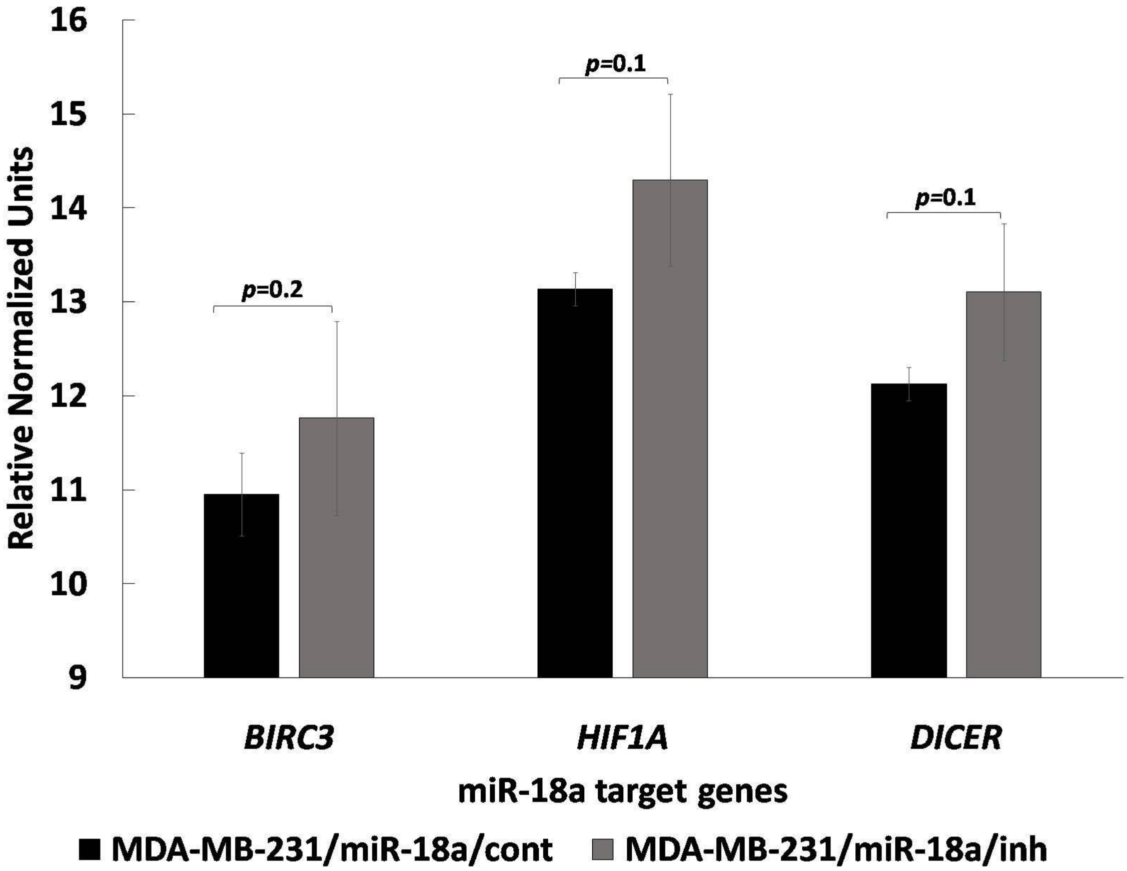

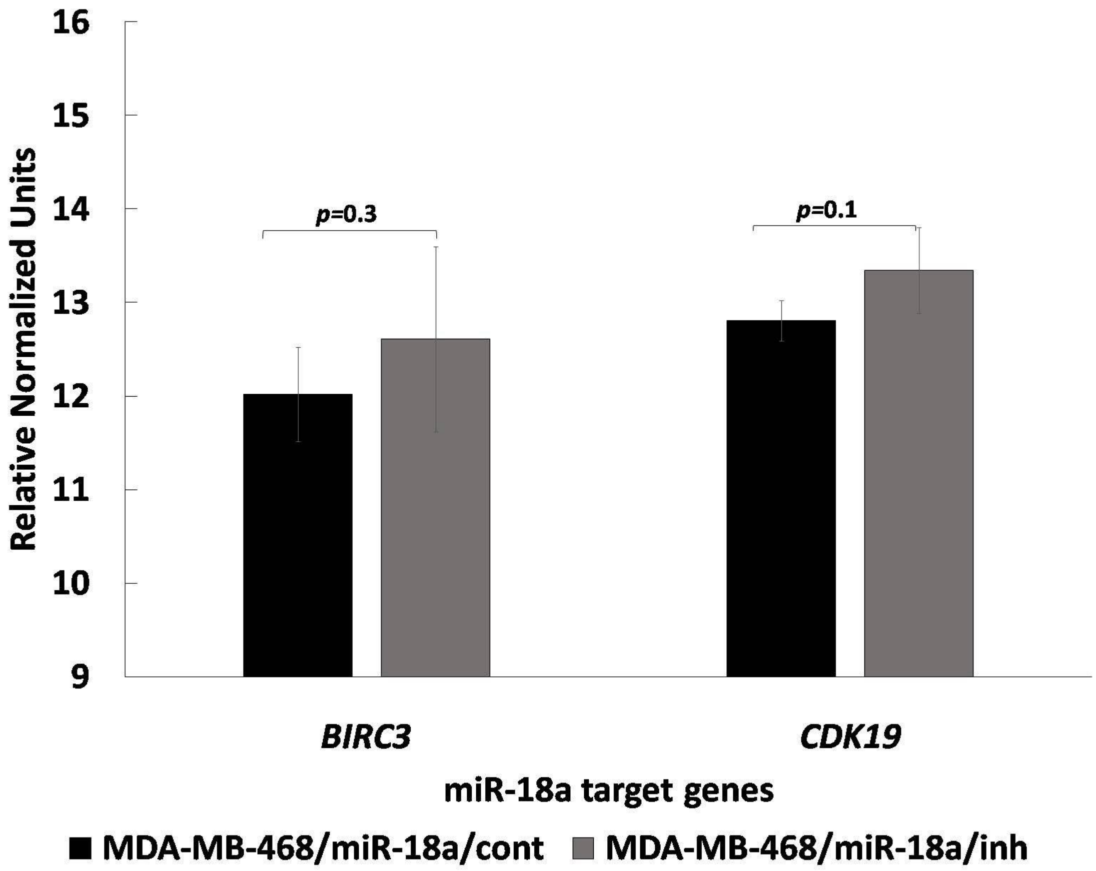

16
15
14
13
12
11
10
9
Relative Normalized Units
p=0.3
p=0.1
BIRC3
CDK19
miR-18a target genes
MDA-MB-468/miR-18a/cont
MDA-MB-468/miR-18a/inh

Supplementary Figure 2: DEGs associated with EMT and drug resistance in miR-18a/low, ER-negative tumours of METABRIC

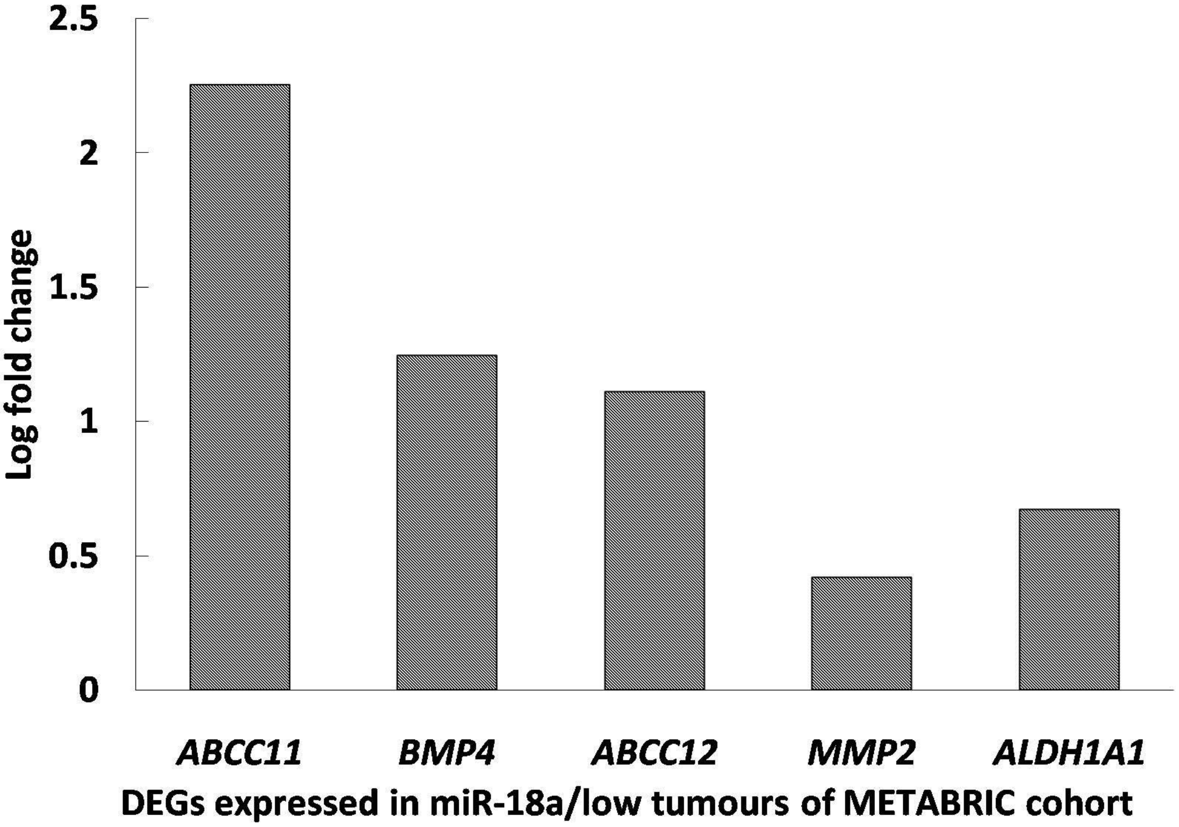

**Supplementary Figure 3: Expression level of stemness genes in ALDH1A1-DsRed2N1 cells vs DsRed2N1 cells.**

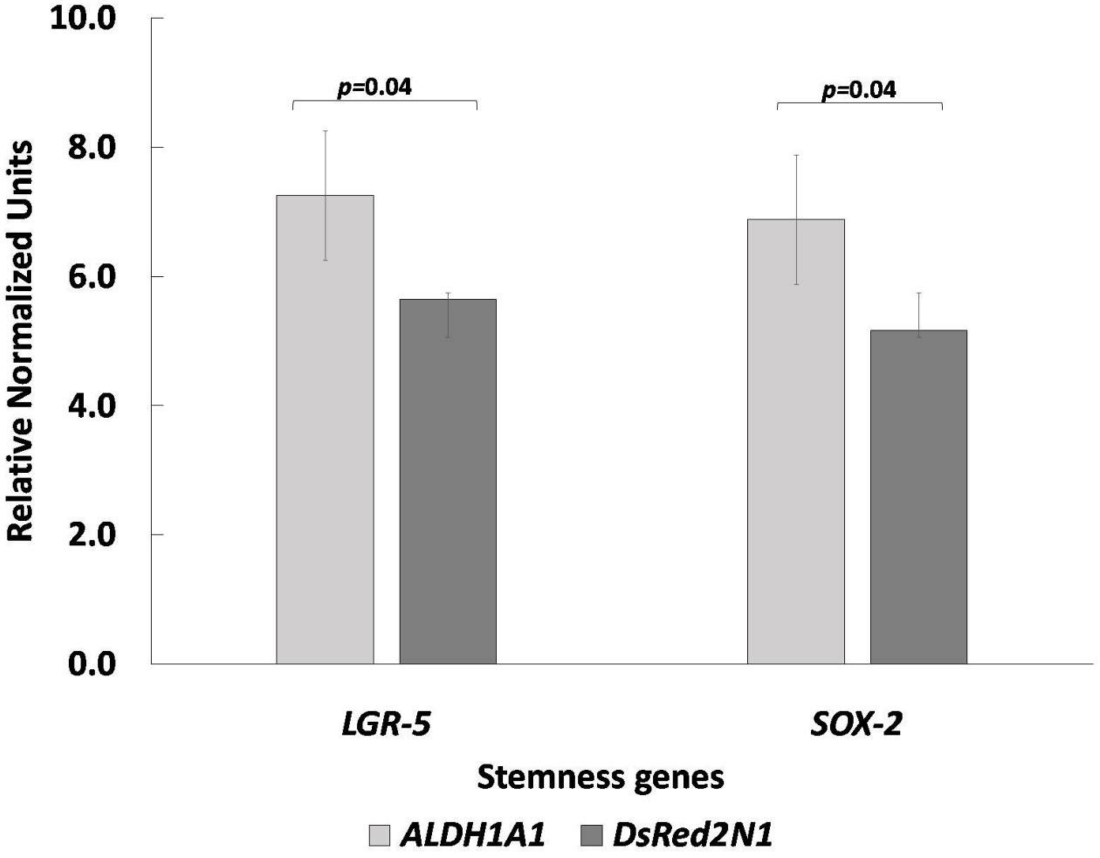

**Supplementary figure 4**: **The heatmap representing the pattern of expression of filtered hypoxia genes in the miR-18a/low tumors of TCGA**

Heatmap depicting expression of filtered hypoxia genes in mir-18a low tumours in TCGA cohort

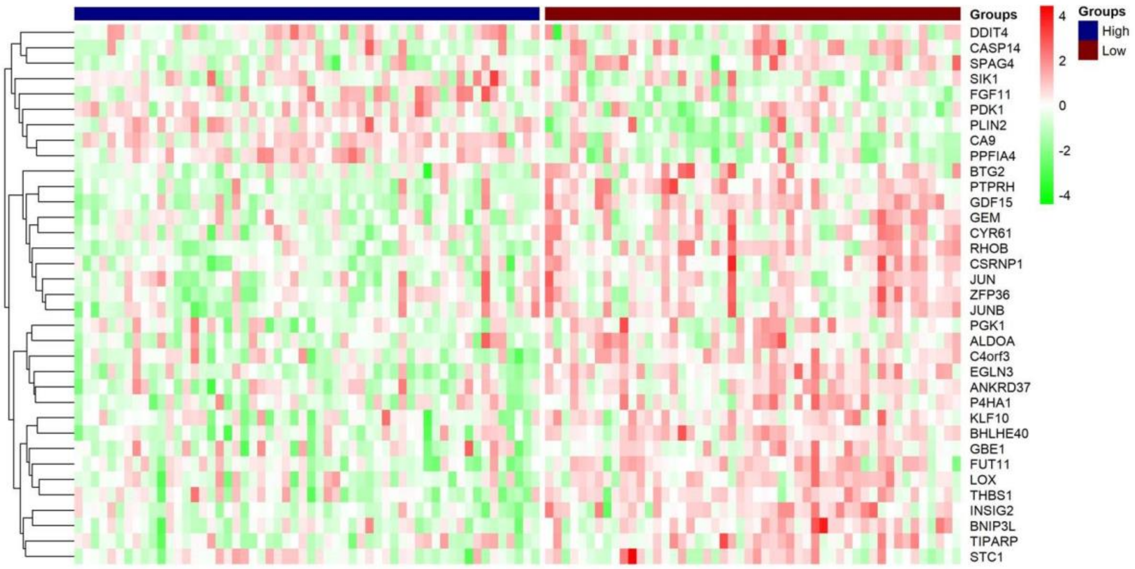

**Supplementary Table S1 : List of antibodies and their dilutions**

| Sl. No | Antibody | Purpose | Product details | Dilution |
|---|---|---|---|---|
| 1 | TNFAIP3 | Western Blot | Rabbit monoclonal EPR2663-ab92324 Abcam | 1:1000 |
| 2 | CD49f | Western Blot | Rabbit monoclonal EPR5578-ab124924 Abcam | 1:5000 |
| 3 | Cytokeratin-19 | Western Blot | Rabbit monoclonal EP1580Y-ab52625 Abcam | 1:10000 |
| 4 | MMP9 | Western Blot | Rabbit monoclonal- PA5-27191 Invitrogen | 1:1000 |
| 5 | RAC3 | Western Blot | Rabbit monoclonal EPR6679(B)-ab124943 Abcam | 1:5000 |
| 6 | ALDH1 | Western Blot | Mouse monoclonal- 611194 BD biosciences | 1:500 |
| 7 | HIF1-α | Western Blot | Rabbit monoclonal EP1215Y-ab51608 Abcam | 1:500 |
| 8 | CD44 | Flow cytometry | CD44 (156-3C11) Mouse mAb-3570 Cell signalling technology | 1:100 |
| 9 | CD24 | Flow cytometry | CD24 (M1/69) Rat mAb PE Conjugate- 90378 Cell signalling technology | 1:660 |
| 10 | E-cadherin | Western Blot /Immunofluorescence | Rabbit monoclonal EP700Y-ab40772 Abcam | 1:5000/1:500 |
| 11 | Vimentin | Immunofluorescence | Mouse clone V9- AM074-5M Bio-genex | 1:25 |
| 12 | Integrin β3 | Immunohistochemistry | Rabbit monoclonal EPR2417Y-ab75872 Abcam | 1:500 |

**Supplementary Table S2. Clinico-pathological characteristics of 211 ER-negative patients: The clinical pathological characteristics of the tumors used for analysis from TCGA tumors**

| | All N (%) (n = 211) | miR-18a/low (n = 50) | miR-18a/high (n = 57) |
|---|---|---|---|
| **Age (y)** | | | |
| Mean | 55 | 57 | 52 |
| Median | 55 | 58 | 51 |
| **Tumor Size (cm)** | | | |
| T1 | 33 (22) | 9 (26) | 4 (10) |
| T2 | 100 (66) | 19 (56) | 34 (83) |
| T3 | 13 (9) | 3 (9) | 2 (5) |
| T4 | 5 (3) | 3 (9) | 1 (2) |
| **Stage** | | | |
| 1 | 20 (14) | 3 (9) | 2 (5) |
| 2 | 96 (65) | 20 (63) | 31 (76) |
| 3 | 29 (20) | 9 (28) | 8 (19) |
| 4 | 2 (1) | | |
| **Lymph Node status** | | | |
| Positive | 71 (39) | 17 (49) | 15 (30) |
| Negative | 112 (61) | 18 (51) | 35 (70) |
| **Menopausal status** | | | |
| Pre | 46 (24) | 12 (26) | 18 (36) |
| Post | 142 (76) | 35 (74) | 32 (64) |
| **Her2 positivity** | | | |
| Negative | 111 (62) | 17 (40) | 39 (76) |
| Positive | 37 (21) | 18 (43) | 4 (8) |
| Equivocal | 31 (17) | 7 (17) | 8 (16) |

**Supplementary Table S3. Clinico-pathological characteristics of 265 ER-negative patients: The clinical pathological characteristics of the tumors used for analysis from METABRIC tumors**

| | All N (%) (n = 265) | miR-18a/low (n = 54) | miR-18a/high (n = 62) |
|---|---|---|---|
| **Age (y)** | | | |
| Mean | 55 | 59 | 53 |
| Median | 55 | 60 | 52 |
| **Tumor Size (cm)** | | | |
| Mean | 2.9 | 2.4 | 2.9 |
| Median | 2.5 | 2.2 | 2.5 |
| **Stage** | | | |
| 0 | 2 (1) | 1 (2) | 0 (0) |
| 1 | 63 (27) | 17 (34) | 9 (15) |
| 2 | 127 (55) | 22 (44) | 42 (71) |
| 3 | 39 (17) | 10 (20) | 8 (14) |
| **Grade** | | | |
| I | 2 (1) | 1 (2) | 0 (0) |
| II | 33 (13) | 11 (22) | 3 (5) |
| III | 223 (86) | 38 (76) | 58 (95) |
| **Lymph Node status** | | | |
| Positive | 139 (52) | 27 (50) | 33 (53) |
| Negative | 126 (48) | 27 (50) | 29 (47) |
| **Menopausal status** | | | |
| Pre | 99 (37) | 13 (24) | 27 (44) |
| Post | 166 (63) | 41 (76) | 35 (56) |
| **Her2 positivity** | | | |
| Negative | 187 (71) | 32 (59) | 59 (95) |
| Positive | 78 (29) | 22 (41) | 3 (5) |

**Supplementary Table S4: List of luminal and basal genes**

| Luminal genes | Basal genes |
|---|---|
| *ESR1* | *CK5/6* |
| *GATA3* | *Laminin* |
| *TFF3* | *CENPI* |
| *FOXA1* | *CENPK* |
| *LIV-1* | *CDC7* |
| *KRT8* | *KIF18A* |
| *GRM4* | *CCNE2* |
| *GRM8* | *STIL* |
| *KRT18* | *CDCA7* |
| *PGR* | *CKS2* |
| *NMUR1* | *MIA* |
| *MUC1* | *ANLN* |
| *CX3CL1* | *FABP7* |
| *NCAM1* | *KRT17* |
| *XBP1* | *KRT6b* |
| | *DCS2* |

**Supplementary Table S5. Clinico-pathological characteristics of 105 ER-negative patients: The clinical pathological characteristics of the tumors used for analysis from our case series.**

| | All N (%) (n= 105 patients) |
|---|---|
| **Age (y)** | |
| Mean | 54 |
| Median | 54 |
| **Tumor Size (cm)** | |
| Mean | 3.5 |
| Median | 3 |
| **Stage** | |
| I | 16 (15) |
| II | 53 (50) |
| III | 32 (30) |
| IV | 4 (4) |
| **Grade** | |
| I | 6 (6) |
| II | 38 (39) |
| III | 54 (55) |
| **Lymph Node status** | |
| Positive | 54 (51) |
| Negative | 49 (47) |
| Nx | 2 (2) |
| **Menopausal status** | |
| Pre | 28(27) |
| Post | 77(73) |
| **Her2 positivity** | |
| Equivocal | 4 (4) |
| Negative | 72 (68) |
| Positive | 29 (28) |

**Supplementary Table S6. Clinical characteristics of post NACT residual tumors: Clinico-pathological characteristics of (n=54) post NACT residual tumors and (n=43) tumors with adequate tissue available for estimation of integrin β3 and miR-18a**

| | n = 54 patients (%) | n = 43 patients (%) |
|---|---|---|
| **Age** | | |
| Mean (Yrs) | 49 | 49 |
| Median (Yrs) | 48 | 48 |
| **Tumor Size** | | |
| Mean (cm) | 6 | 6 |
| Median (cm) | 6 | 6 |
| **Stage** | | |
| III | 38 (70) | 31 (72) |
| IV | 14 (26) | 10 (23) |
| Nx | 2 (4) | 2 (5) |
| **Menopausal status** | | |
| Pre | 24 (44) | 19 (44) |
| Post | 30 (56) | 24 (56) |
| **Estrogen Receptor** | | |
| Positive | 30 (56) | 22 (51) |
| Negative | 24 (44) | 21 (49) |
| **Progesterone Receptor** | | |
| Positive | 28 (52) | 21 (49) |
| Negative | 26 (48) | 22 (51) |
| **HER2** | | |
| Positive | 18 (33) | 13 (30) |
| Negative | 29 (54) | 25 (58) |
| Equivocal | 7 (13) | 5 (12) |

**Supplementary Table S7: List of primer sequences used in qPCR**

| Sl No. | Gene | PRIMER SEQUENCE |
|---|---|---|
| 1 | *ACTB* | 5’-TTCCTGGGCATGGAGTC-3’<br>3’-CAGGTCTTTGCGGATGTC-5’ |
| 2 | *ANLN* | 5’-ACAGCCACTTTCAGAAGCAAG-3’<br>3’-CGATGGTTTTGTACAAGATTTCTC-5’ |
| 3 | *BCL2* | 5’-TACCTGAACCGGCACCTG-3’<br>3’-GCCGTACAGTTCCACAAAGG-5’ |
| 4 | *BIRC3* | 5’-CCATGGGTTCAACATGCCAAGTGGT-3’<br>3’-GGGTAACTGGCTTGAACTTGACGG-5’ |
| 5 | *BMPR1B* | 5’-ATTCCCAAACCGGTGGAGCAGT-3’<br>3’-TTGATGCAGGATTGTGAGCCCAGC-5’ |
| 6 | *CDK19* | 5’-TACCTCCATGCAAATTGGGTGCT-3’<br>3’-R-TTTGACTCTCCCCCTCTCAGGA-5’ |
| 7 | *DICER* | 5’-TTAACCTTTTGGTGTTTGATGAGTGT-3’<br>3’-GCGAGGACATGATGGACAATT-5’ |
| 8 | *HIF1A* | 5’-TGCTTACACACAGAAATGGCCT-3’<br>3’-TAGTTAGGGTACACTTCATTCTGAG-5’ |
| 9 | *ITGB3* | 5’-CCCACCAGAGGCCCTCGAAA-3’<br>3’-AAGCGGGTCACCTGGTCAGT-5’ |
| 10 | *LGR5* | 5’-GACCATTGCCTACACCAAGC-3’<br>3’-GAGCAACAGGGCAATGTGTT-5’ |
| 11 | *PGR* | 5’-TTATAATTCGAGGCGGTTAGTGTTT-3’<br>3’-TCGAACTTCTACTAACTCCGTACTACGA-5’ |
| 12 | *PUM1* | 5’-CCGGAGATTGCTGGACATATAA-3’<br>3’-TGGCACGCTCCAGTTTC-5’ |
| 13 | *RPLP0* | 5’-GGCTGTGGTGCTGATGGGCAAGAA-3’<br>3’-TTCCCCCGGATATGAGGCAGCAGT-5’ |
| 14 | *SALL4* | 5’-GGCCAATAGTCAAGCCGAAA-3’<br>3’-TCCGACCTTCCATCTCAGTG-5’ |
| 15 | *ZEB2* | 5’-TGCACAGAGTGTGGCAAGGC-3’<br>3’-TGGGCACTCGTAAGGTTTTTCACC-5’ |
